# Sensitivity of Pulmonary 4D Flow MRI Biomarkers to Controlled Spatial and Temporal Degradation in a CFD Benchmark

**DOI:** 10.64898/2026.09.30.755784

**Authors:** Aitor Zubillaga-Unsain, Cesar Caballero-Gaudes, Jesús Ruiz-Cabello

## Abstract

**Purpose:** To quantify, in a computational fluid dynamics (CFD) benchmark, how spatial degradation, temporal acquisition and noise change the error of plane-based pulmonary-artery flow endpoints.

**Methods:** Twenty-two historical, non-converged CFD simulations (11 geometry families) were voxelised at 2 mm with 80 frames per cycle. Simulations with numerical divergence were excluded, leaving 13 (7 families). Fields were degraded to 4 and 8mm and with noise, plane waveforms were sampled at 20 phases with one-interval averaging, and errors were computed against the noise-free reference waveform on 66 geometric planes. Family-mean paired differences are reported with t-based 95% confidence intervals under an analysis plan fixed in advance.

**Results:** Spatial degradation from 2 to 4mm increased full-waveform nRMSE by 0.0032 [0.0021, 0.0043], in 7 of 7 families; cycle-mean flow bias accounted for 23% of the squared error, against 60% at 8 mm. With SNR scaled to voxel volume, the increase was not resolved (0.0002 [−0.0012, 0.0015]). Twenty-phase acquisition increased PI error (0.0223 [0.0214, 0.0233]) and diastolic-trough timing error (0.63 [0.52, 0.74] frames) more than spatial degradation, and its point estimates for these endpoints were larger with every recovery method; the nRMSE ordering and peak timing depended on the recovery method. The implementation of the downsampling operator mattered: an alternative, corner-aligned implementation of the same 4mm degradation gave a larger spatial effect (nRMSE difference 0.0010 [0.0002, 0.0017]).

**Conclusion:** In this benchmark, a 2 to 4mm degradation produced a small, consistent, mostly shape-related nRMSE increase that depended on the noise model, whereas 20-phase acquisition changed PI error and trough timing more. The results describe benchmark behaviour, not clinical acquisition requirements.

## 1 Introduction

Pulmonary 4D flow MRI provides hemodynamic measures that are studied in pulmonary hypertension, including flow waveforms, the pulsatility index (PI), feature timing, and vortex- and wall-derived quantities ^1–5^. These measures are computed from velocity fields of finite spatial and temporal resolution. Consensus statements recommend reporting acquired and reconstructed resolution and assessing its influence ^6,7^. Phantom and CFD studies in the aorta and carotid bifurcation show that sensitivity to resolution depends on the quantity measured ^8–10^. Work on view sharing distinguishes the nominal number of cardiac phases from the effective temporal averaging window ^11–13^. We are not aware of equivalent controlled data for pulmonary-artery plane-flow endpoints.

Clinical pulmonary 4D flow protocols balance voxel size, phase count and scan time; the cohort from which the geometries of this study were derived was acquired at 2.5mm isotropic resolution with 20 cardiac phases ^16^. Which of these choices matters depends on the endpoint. The cycle-mean flow, the shape of the waveform, the pulsatility index and the timing of the systolic peak and diastolic trough respond to different parts of the acquisition, and an error in one need not appear in another. Separating these effects requires a reference that is known on every plane and at every frame, which in vivo data cannot provide. Velocity fields from CFD, voxelised on a fine grid, can serve as such a reference, provided that their own numerical limitations are stated with the results.

Our previous synthetic pulmonary-artery study found that voxel-level reconstruction fidelity did not rank PI accuracy among near-equivalent super-resolution models ^14^. The present study addresses a separate question: a deterministic benchmark that degrades CFD-derived reference fields along controlled axes and measures how the error of plane-flow endpoints responds. We compare a 2 to 4mm spatial degradation with 20-phase acquisition on four plane-flow endpoints: full-waveform nRMSE, PI error, and systolic-peak and diastolic-trough timing. We split the waveform error into cycle-mean flow bias and waveform shape, and ask how the comparison depends on the recovery method, the noise model and the degradation operator. The study does not simulate a scanner acquisition and makes no acquisition recommendation.

## 2 Methods

### 2.1 Study design

Twenty-two CFD cases were analysed: 11 pulmonary-artery geometries with two flow simulations each (Supplementary Table S1), one with the SST *k*–*ω* turbulence model and one laminar, with the same boundary conditions and solver settings apart from the exceptions listed in Supplementary Note S2. A geometry and its two flow simulations form a family, the unit of inference; cases, planes, sampling offsets, recovery methods, spatial paths and noise realisations are repeated measures within a family.

The analysis plan and the case-exclusion rule were fixed before any result of this analysis was read; their design was informed by earlier internal analyses of the same simulations (Supplementary Note S8).

### 2.2 Reference data

The eleven geometries were derived from cardiac magnetic resonance (CMR) examinations of patients with group 2 pulmonary hypertension enrolled in the SAPPHIRE study ^15,16^, a single-centre prospective observational study (Hospital Universitario 12 de Octubre and CNIC, Madrid, Spain) in which patients underwent right heart catheterisation and CMR, including 4D flow MRI, within 24 h. The examinations used here are identified internally as HPIC01–HPIC19 and belong to the SAPPHIRE cohort. The pulmonary-artery lumen was manually segmented from 11 of the 19 examinations. Only the vessel anatomy of each patient was used; apart from the common inlet waveform described below, no patient flow or clinical data entered the benchmark. The transient simulations (Ansys Fluent 2026 R1) used meshes of 0.76–0.98 million cells (equivalent cell edge 0.42–0.54mm), and one cardiac cycle of each was exported as 80 frames of 12.5 ms per 1.0 s cycle (solver settings in Supplementary Note S2). The historical simulations were not numerically converged; we use them as realistic flow fields rather than as converged solutions, and exclude cases with numerical divergence (Section 2.3). The prescribed inlet speed was spatially uniform over the dome-shaped inlet surface and applied along the local surface normal. Its temporal variation followed a five-harmonic Fourier-series waveform derived from the flow waveform of one patient and was normally scaled by a factor of 0.2; the two run-specific exceptions are described in Supplementary Note S2. All outlets were at 0 Pa gauge pressure, with reverse flow allowed and the backflow direction normal to the outlet. Blood was modelled as a Newtonian fluid (density 1050 kg/m^3^, viscosity 0.0035 Pa s), and the walls as rigid and no-slip.

The cell-centre velocities of the exported frames were averaged, without volume weighting, into 2mm isotropic voxels. The vessel mask marks voxels that contain at least one cell centre, and voxels outside it are zero. Empty voxels inside the closed vessel surface and away from the open caps (111 voxels in 10 cases) were filled by inverse-distance weighting of the nearest cells. On every plane, the noise-free reference plane waveform is the plane integral of this undegraded field, and every endpoint error is computed against this noise-free reference waveform. It is a property of the voxelised field, not a solver boundary flux or the flux through the true lumen section.

### 2.3 Case exclusion

Cases were classified once, before any benchmark result was read, by a written rule that uses only the quality of the exported CFD fields (Table 1; Supplementary Tables S6 and S7). The rule tests the speed outside the caps (at most 3 m/s beyond 16mm from any open cap), numerical divergence (no cell above 10 m/s and no sustained growth of the maximum speed at the end of the cycle), flow conservation between the inlet and mid-trunk sections (within 5%), inlet artefacts reaching the inlet section, high-speed cells near the measurement sections, and start-up transients (an exported cycle that overlaps a documented inlet ramp), always on the raw, unrepaired fields. One case, whose exported flow corresponds to five times the inflow of the others, was excluded explicitly. The primary cohort contains the cases that fail none of these tests, and so the cases without detectable divergence in their exported fields; a broad sensitivity cohort omits the divergence and start-up tests. Part of the rule was written after the CFD-quality values had been computed (Supplementary Note S7).

**Table 1.** Case cohorts. The exclusion tests are described in the Methods; criteria and per-case outcomes are in Supplementary Tables S6 and S7.

| Cohort | Exclusion tests applied | Cases | Families | Use |
| --- | --- | --- | --- | --- |
| Primary<br>(non-divergent) | speed outside the caps, divergence, flow conservation, inlet artefacts, cap-zone outliers, start-up transients | 13 | 7 | primary analysis |
| Broad | speed outside the caps, flow conservation, inlet artefacts, cap-zone outliers; no divergence or start-up test | 19 | 10 | sensitivity analysis |

### 2.4 Measurement planes

Planes were placed by a geometric rule on the exact source surface of each case (wall and labelled open caps); no velocity, flow or error value was used. From every open cap, the lumen was tracked inward by cross-sections at 0.5mm steps, and a station was eligible only if:

- (P1) the integration disk lay at least 16mm from every open-cap vertex, with no cap crossing;
- (P2) no other lumen lay within the disk radius plus 4mm;
- (P3) the section loop was closed by wall triangles only;
- (P4) the plane normal deviated at most 20° from the local centreline;
- (P5) the equivalent diameter was at least 12mm;
- (P6) the station lay at least 0.5 equivalent diameters upstream of the bifurcation;
- (P7) the ratio of maximum to equivalent radius was at most 1.5;
- (P8) the voxelised section, sampled as the integrator samples it, formed a single region inside the disk, ignoring regions that lie entirely outside the closed vessel surface.

The disk radius was the maximal section radius plus 8mm. The pre-declared locations were the first eligible trunk station from the inlet, the station nearest the midpoint between the inlet station and the bifurcation, and the first eligible station from each outlet. 66 of 88 candidate measurements were valid (Figure 1), including the inlet of every case.

**Figure 1.**
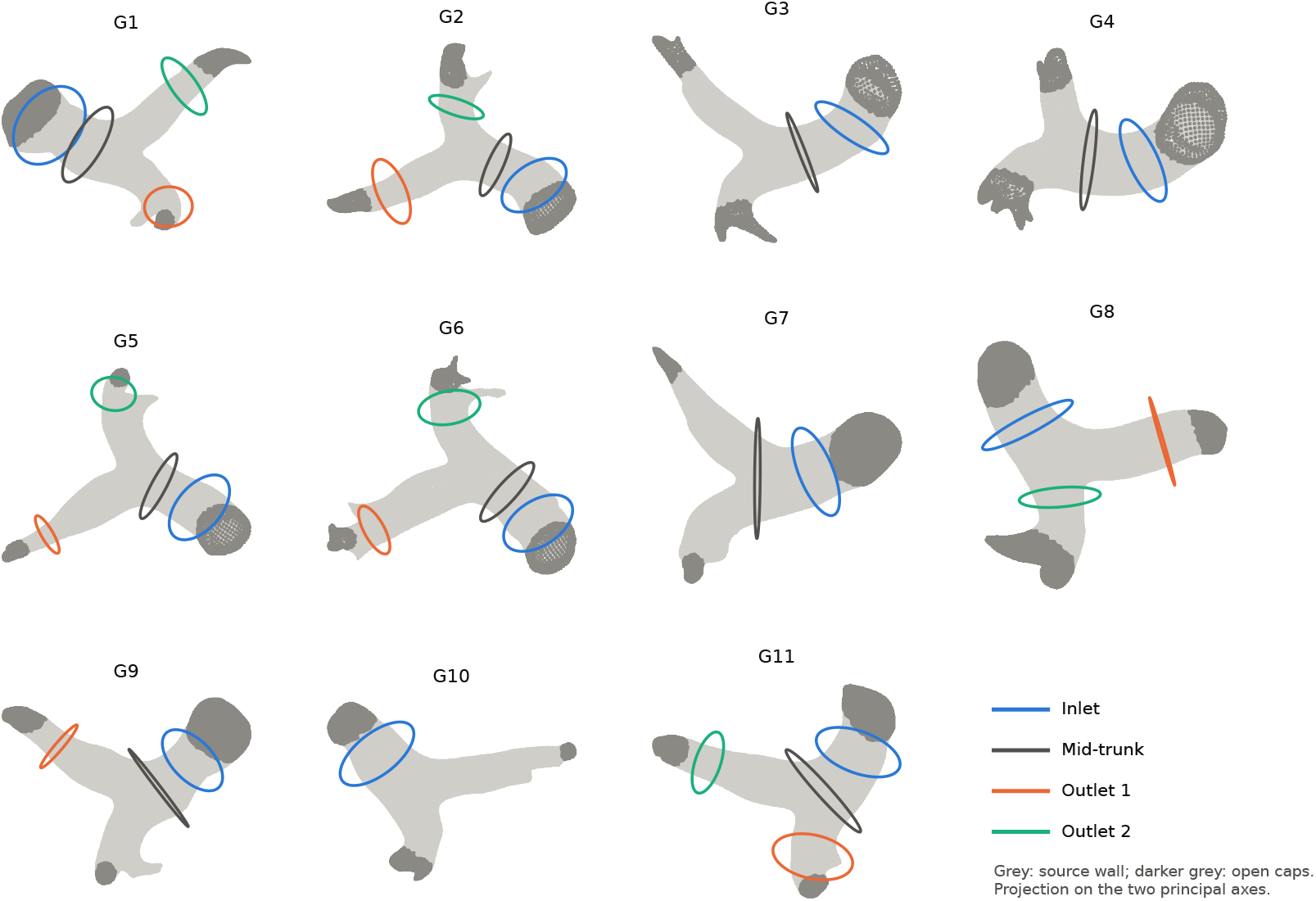
Measurement planes on the source surfaces of the 11 geometries (G1–G11; first flow simulation shown), projected on each vessel’s two principal axes. Circles are integration disks of accepted planes.

### 2.5 Degradations and recovery

The design is summarised in Table 2 and Figure 2. The clinical 4D flow protocol of the source cohort used 2.5mm isotropic resolution and 20 cardiac phases ^16^; the 2 to 4mm spatial contrast brackets its spatial resolution, and the 20-phase level matches its temporal sampling.

**Table 2.** Degradation design.

| Factor | Levels |
| --- | --- |
| Baseline condition | 2 mm, 80 frames per cycle, point sampling, 27.5 dB noise (mean of 10 realisations); every contrast is a change in error relative to it |
| Reference waveform | plane integral of the undegraded, noise-free 2 mm field on the same plane; every error is computed against it |
| Spatial levels | 2 mm (reference); 4 mm (primary) and 8 mm: Gaussian point-spread function ($\sigma = 1.6$ and 3.2 mm) sampled at block centres; 4 mm with the earlier operator (sensitivity) |
| Temporal acquisition | 80, 40, 32, 25, 20, 16, 10 phases per cycle (20 phases = 50 ms); point, box (one output interval) or triangle (two output intervals) kernels; four sampling offsets |
| Noise | none; fixed per-voxel SNR of 27.5 and 17.5 dB (primary); SNR scaled with voxel volume, 45.56 and 35.56 dB at 4 mm (secondary); 10 realisations |
| Temporal recovery | periodic linear (primary); PCHIP, sinc/Fourier, nearest-sample assignment (sensitivity) |
| Spatial paths | native coarse grid and trilinear interpolation to 2 mm, averaged |
| Measurement planes | 66 of 88 candidate measurements; primary: trunk sections (inlet, mid-trunk); branch sections secondary |
| Cohorts and inference | case-exclusion rule: 13 cases in 7 families (primary), 19 cases in 10 families (broad; Table 1); family-level t-based 95% CI and exact sign-flip test |

**Figure 2.**
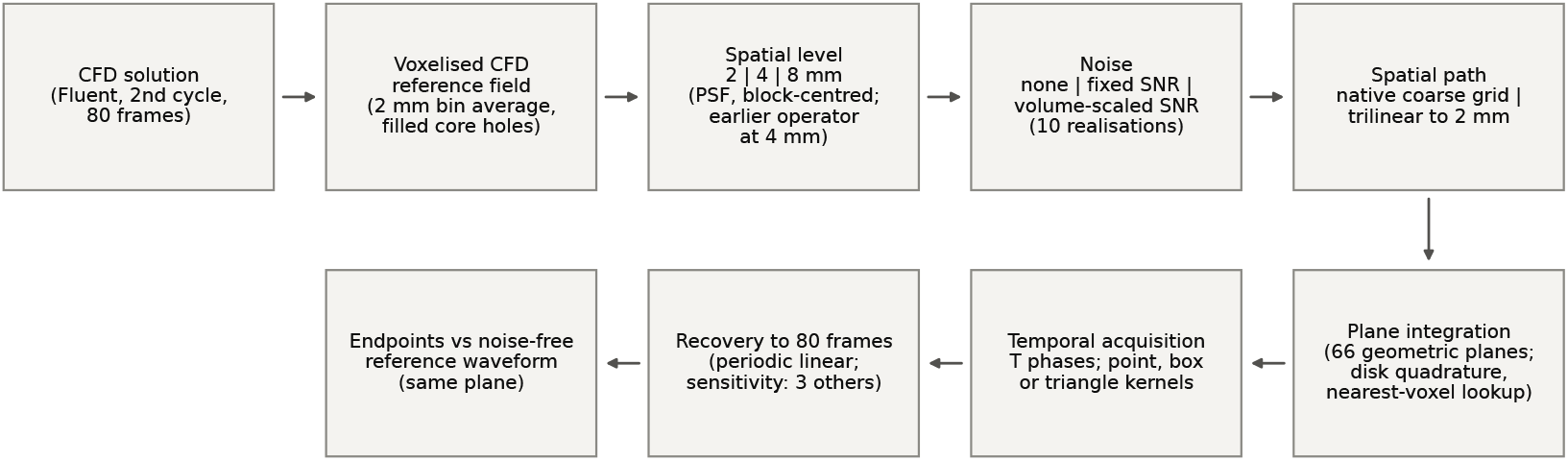
Processing chain. Endpoint errors are computed against the noise-free reference waveform on the same geometrically defined plane.

For spatial degradation, each coarse voxel covers a block of *f*^3^ reference voxels (*f* = 2 at 4mm, 4 at 8mm), and its value is the reference field filtered by a Gaussian point-spread function and sampled at the block centre; we call this the block-centred operator. The point-spread function has *σ* = 0.4*f* reference voxels (1.6mm at 4mm), a full width at half maximum of about 0.94 coarse voxels, which approximates the aperture of a coarse voxel, as in our previous study ^14^.

The degraded field was evaluated on its coarse grid and after trilinear interpolation back to 2mm, and the two results were averaged. The 8mm level is descriptive; it shows how the split between bias and shape changes with a further coarsening, without a test. As a sensitivity analysis, the 4mm level was also produced with the corner-aligned trilinear downsampling of our previous study, which we call the earlier operator: its lattice spans the full reference extent, so its pitch exceeds the nominal voxel size and its samples are displaced by up to about one coarse voxel from the block centres (Supplementary Note S3).

Zero-mean Gaussian noise was added to each velocity component, voxel and frame of the degraded 80-frame field, inside and outside the vessel mask, with standard deviation RMS/10^SNR/20^, where RMS is the in-mask velocity at that spatial level. The primary model holds the per-voxel SNR at 27.5 or 17.5 dB at every level, so that the spatial contrast measures the effect of the operator at equal relative noise. In MRI, larger voxels gain SNR in proportion to their volume at a fixed acquisition time, so a secondary model scales the SNR with voxel volume (45.56 and 35.56 dB at 4mm). Each noisy condition has 10 realisations, and within a spatial level all temporal conditions share the same noise fields. Because noise is added before temporal averaging, averaging and interpolating recovery also reduce the noise in the recovered waveform; since the baseline condition contains noise, operators that smooth noise can reduce error relative to it, and noise-free contrasts are therefore reported as well.

Temporal acquisition acts on the integrated 80-frame plane waveform, which is exact because flux integration commutes with the linear kernels used. Each of the *T* samples per cycle is a kernel-weighted average of the waveform centred on its nominal sample time: a Dirac kernel (point sampling), a box of one output interval, or a triangle of two output intervals (Figure 3; sample times and interpolation in Supplementary Note S2). The kernels are even about the sample time, so the time assigned to each sample is its kernel centroid and no phase shift is introduced; a kernel that is not centred on its sample time would shift every timing endpoint. The box and triangle kernels tile the cycle and preserve the cycle-mean flow. Four sampling offsets were used for *T* < 80.

**Figure 3.**
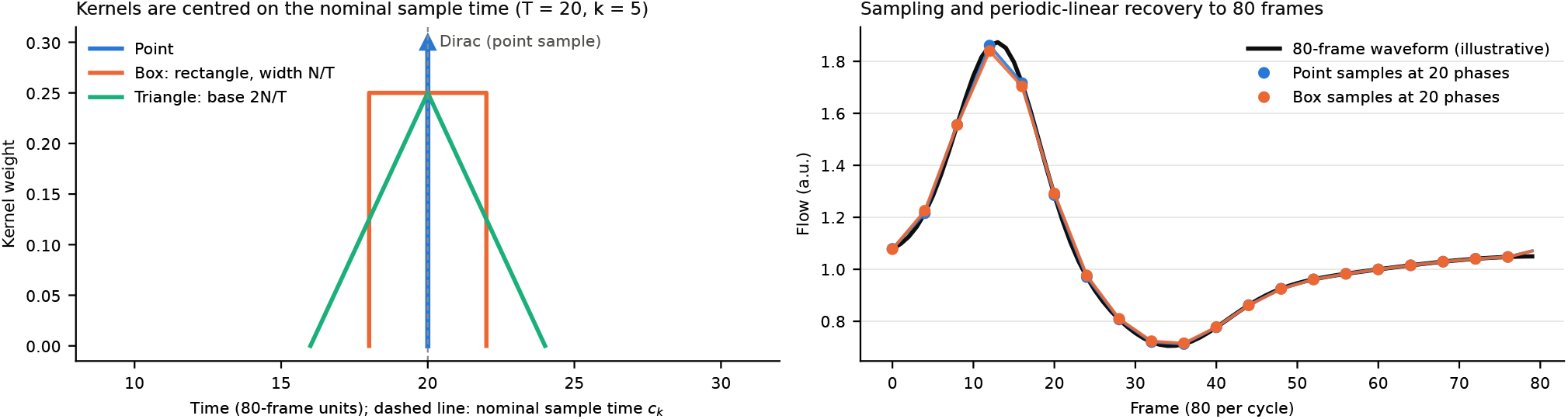
Temporal acquisition operator. Left: point, box and triangle kernels centred on the nominal sample time *c*_*k*_ (*T* = 20). Right: illustrative waveform sampled at 20 phases with the point and box kernels and recovered to 80 frames by periodic-linear interpolation.

Through-plane flow was integrated over the whole integration disk, without a lumen mask (quadrature in Supplementary Note S2). In the reference field the voxels outside the vessel are zero; after degradation, the disk integral also includes the velocity that the point-spread function spreads beyond the lumen, and the noise in the margin of the disk outside the lumen. Integrating without a mask captures the flow displaced outside the lumen by blurring and avoids a lumen mask whose definition would itself depend on the resolution. Waveforms sampled at *T* < 80 were recovered to 80 frames by periodic-linear interpolation (primary) or by PCHIP (piecewise cubic Hermite), sinc/Fourier or nearest-sample assignment (sensitivity). Linear and PCHIP recovery place recovered extrema at sample times and so give identical timing endpoints.

### 2.6 Endpoints

The primary endpoint was full-waveform nRMSE, RMSE(*Q*_pred_ − *Q*_ref_ )/(*P*_95_ − *P*_5_) over the 80 frames, where *P*_95_ − *P*_5_ is the pulsatile range of the reference waveform; it contains both the cycle-mean flow bias and the waveform-shape error. The headline endpoints were the absolute error of the flow pulsatility index, PI = (*Q*_sys_ − *Q*_dia_)/|*Q*_mean_|, and the unsigned periodic timing errors of the systolic peak and the diastolic trough, in frames of 12.5 ms. With *d* = *Q*_pred_ −*Q*_ref_, the squared nRMSE splits exactly into the squared cycle-mean flow bias and the squared mean-removed nRMSE; we report the signed change in cycle-mean flow, the mean-removed nRMSE and the bias share of the squared error. Small floors on both denominators, derived from the reference waveforms, were never active (Supplementary Note S2).

### 2.7 Contrasts and statistics

Each contrast is the change in endpoint error relative to the baseline condition: 2mm, 80 frames, point sampling at offset 0 and 27.5 dB noise. The baseline condition contains noise so that every contrast compares two noisy measurements, as an acquisition study would; the noise-free contrasts isolate the operators themselves. The primary contrast is spatial degradation from 2 to 4mm at 80 frames. The others are 20-phase acquisition (20 phases with one-interval box averaging, which changes the phase count, the per-phase averaging and the effective noise together), the averaging footprint (triangle vs box at 20 phases), noise (17.5 vs 27.5 dB), combined 4mm and 20-phase acquisition, and degradation to 8mm. The spatial and 20-phase contrasts were compared directly, as paired within-family differences, on full-waveform nRMSE, PI error, trough and peak timing, and mean-removed nRMSE. Because voxelisation inflates the flux through the smaller branch sections, the primary estimate uses the trunk. For each case, the test-minus-baseline difference was averaged over the valid trunk sections (inlet and mid-trunk), the two spatial paths, the four offsets and the noise realisations; case values were averaged within family, and families with equal weight.

Each estimate is the mean of the family values, with a t-based 95% confidence interval (CI) and an exact two-sided sign-flip test over all sign assignments; a family bootstrap gives a secondary interval (Supplementary Table S15). With 7 families the smallest attainable sign-flip p value is 0.016 (0.002 with the 10 families of the broad cohort). The secondary analyses were the branch sections, noise-free contrasts, each spatial path separately and the volume-scaled noise model. The sensitivity analyses were the three alternative recovery methods, the inlet sections only, a linear drift correction of the non-periodic reference cycles (Supplementary Note S2), the broad cohort without one case whose exported cycle lies under an inlet ramp, and the earlier operator. Two further analyses are labelled post hoc: a split of the 20-phase contrast into its phase-count and averaging parts, and the direction of the PI change.

## 3 Results

Estimates are family means of paired differences with t-based 95% CIs and exact sign-flip p values; CIs and p values are unadjusted for multiplicity. We describe one effect as larger than another only when the 95% CI of their difference excludes zero, and an effect whose interval includes zero as not resolved. The smallest standardised effect detectable with 80% power is 1.27 between-family standard deviations (1.00 in the broad cohort). Full results for both cohorts, family-bootstrap intervals and the few summaries in which the interval and the p value disagree are given in Supplementary Note S9.

### 3.1 Cohorts

The exclusion rule retained 13 cases in 7 families for the primary cohort and 19 cases in 10 families for the broad cohort (Table 1). Eight cases of four geometries showed numerical divergence at the inlet cap and were excluded from the primary cohort; the two cases of one of these geometries also failed the speed test and are absent from the broad cohort as well. One case was excluded explicitly, so its geometry enters both cohorts through its other case only.

### 3.2 Spatial degradation

Figure 4 shows both degradations for the inlet section of one case. Doubling the voxel size from 2 to 4mm increased full-waveform nRMSE by 0.0032 [0.0021, 0.0043] at 27.5 dB and by 0.0036 [0.0021, 0.0051] without noise, that is, by about 0.32% of the reference pulsatile range; the increase occurred in every family (p = 0.016; Figure 5). Most of it was waveform-shape error: the cycle-mean flow bias carried 23% of the squared error at 4mm and 60% at 8mm (Figure 6), the mean-removed nRMSE increased by 0.0025 [0.0017, 0.0033], and the signed change in cycle-mean flow, 0.08 [−0.09, 0.24]%, was not resolved. PI error increased by 0.0054 [0.0025, 0.0082] at 27.5 dB and by 0.0056 [0.0028, 0.0083] without noise. Timing errors are counted in frames of 12.5 ms. The intervals of the spatial changes in trough timing (0.04 [−0.04, 0.11] frames) and peak timing (0.14 [−0.01, 0.29] frames) included zero; for 20-phase acquisition, the peak timing result depended on the recovery method (see the sensitivity analyses). The broad cohort agreed, except that its spatial peak timing change was resolved (0.15 [0.05, 0.26] frames).

**Figure 4.**
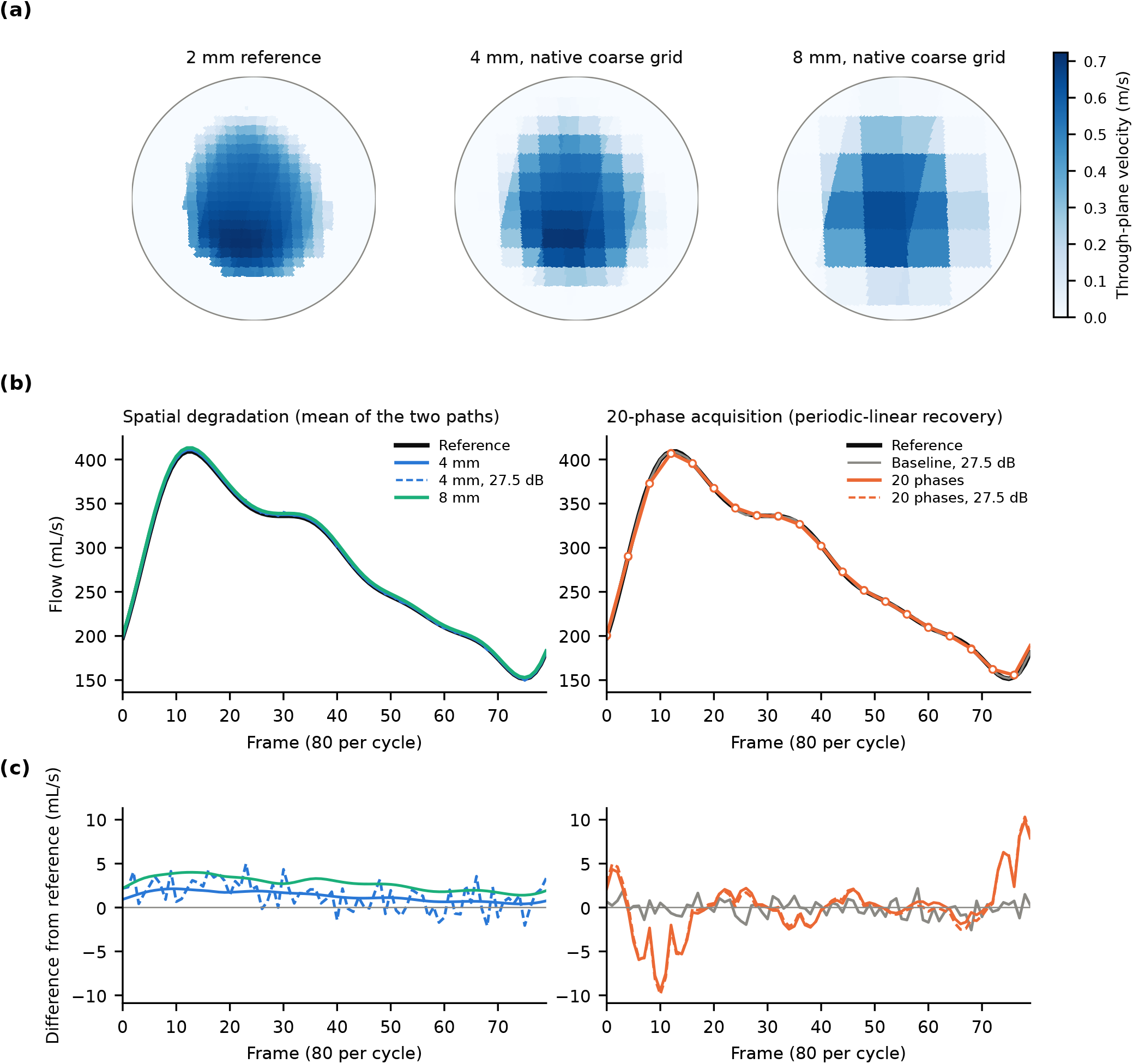
One illustrative case (inlet section of geometry G2, primary cohort). (a) Noise-free through-plane velocity at peak systole at 2mm (reference) and after degradation to 4 and 8mm. (b) Plane flow waveforms under spatial degradation (left) and 20-phase acquisition with box averaging and periodic-linear recovery (right; markers show the 20 samples). (c) Difference from the noise-free reference waveform. Noisy curves show one of the 10 noise realisations. Family-level results are in Figures 5–7.

**Figure 5.**
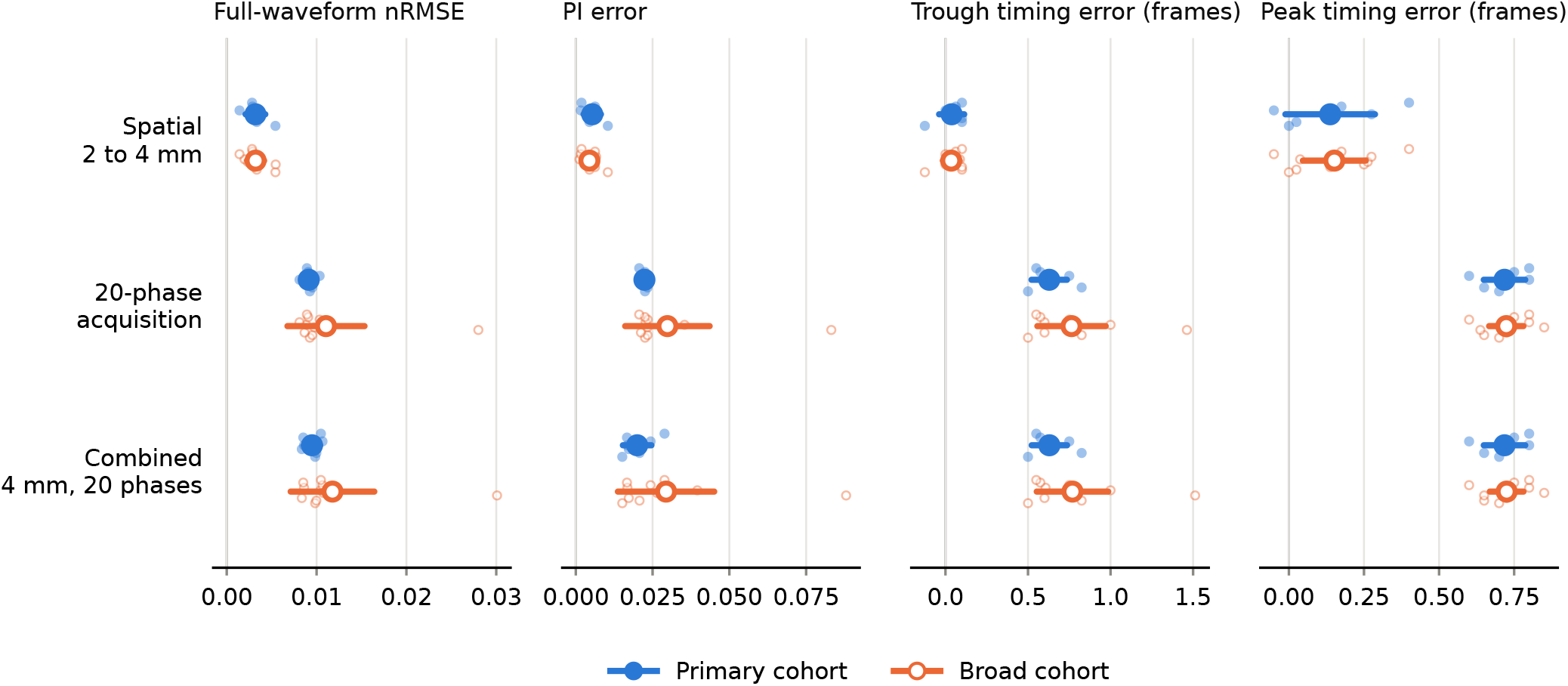
Headline contrasts relative to the baseline condition, with periodic-linear recovery. Large points and bars: family means with t-based 95% CIs; small light points: the values of the individual families. Filled: primary cohort (7 families); open: broad cohort (10 families). The outlying open points in the 20-phase and combined rows (nRMSE, PI and trough timing) belong to geometry G3, which is in the broad cohort only. Timing errors are in frames of 12.5 ms.

**Figure 6.**
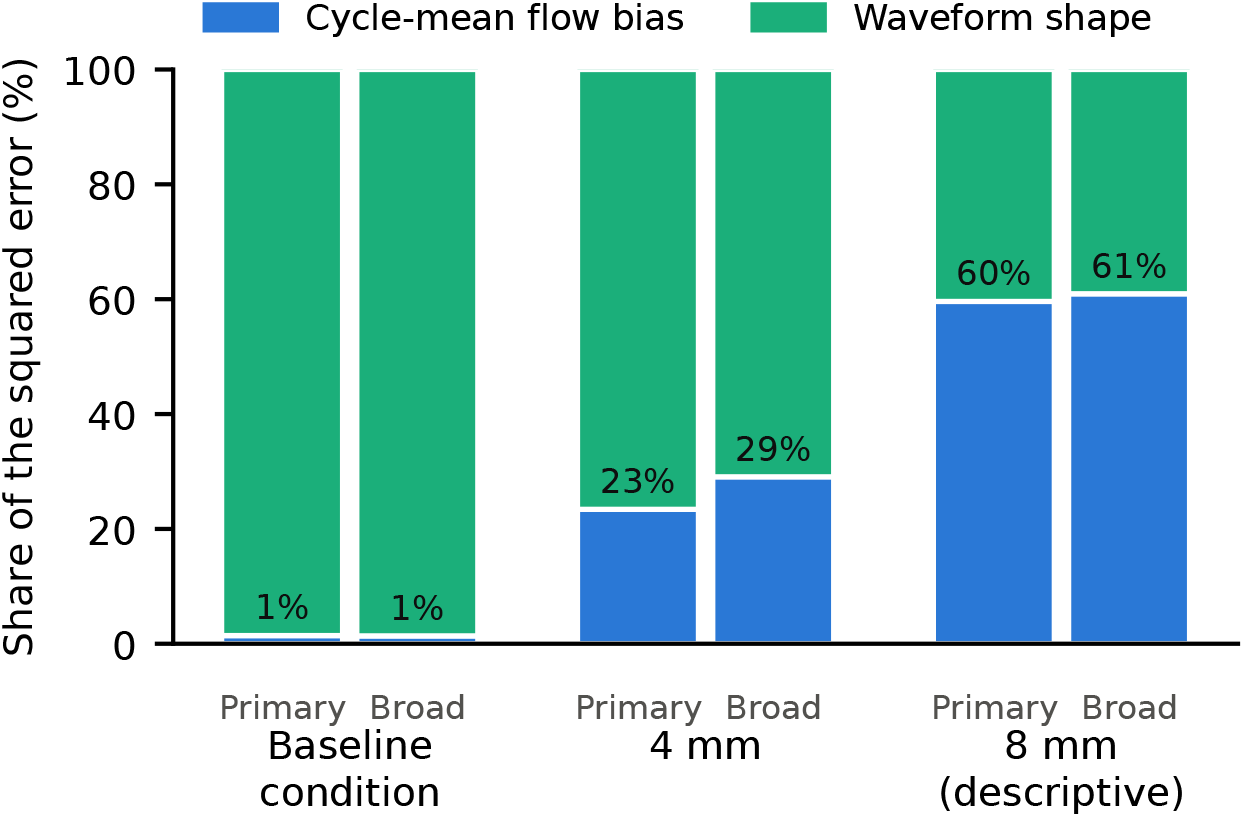
Share of the squared full-waveform error carried by the cycle-mean flow bias and by waveform shape, in the baseline condition and after spatial degradation to 4mm and to 8mm.

### 3.3 Twenty-phase acquisition

Twenty-phase acquisition with one-interval averaging increased full-waveform nRMSE by 0.0091 [0.0085, 0.0098] at 27.5 dB and by 0.0126 [0.0123, 0.0128] without noise, PI error by 0.0223 [0.0214, 0.0233] and 0.0258 [0.0252, 0.0263], trough timing error by 0.63 [0.52, 0.74] frames and peak timing error by 0.72 [0.65, 0.79] frames (Figure 5), and each of these effects exceeded the corresponding spatial effect. The paired differences from the spatial contrast were −0.0059 [−0.0074, −0.0045] for nRMSE and −0.0170 [−0.0199, −0.0141] for PI error (spatial minus 20-phase), and 0.59 [0.48, 0.70] and 0.58 [0.49, 0.67] frames for trough and peak timing (20-phase minus spatial); on mean-removed nRMSE the difference was −0.0066 [−0.0078, −0.0055]. For a noise-free waveform, sampling at 20 phases alone displaces an integer-frame extremum by 1.0 frame on average (sampling-grid quantisation). The 20-phase contrast left the cycle-mean flow unchanged, so its error was entirely waveform shape (bias share 0%). The broad cohort agreed on every comparison. Combining 4mm with 20-phase acquisition gave effects close to those of 20-phase acquisition alone (nRMSE 0.0095 [0.0086, 0.0104]); the footprint and noise contrasts are in Supplementary Table S8.

### 3.4 Secondary and sensitivity analyses

The spatial effect on nRMSE depended on the noise model and on the degradation operator, whereas the 20-phase point estimates for PI error and trough timing stayed above the spatial ones with every recovery method. With the SNR scaled in proportion to voxel volume, the spatial nRMSE increase was 0.0002 [−0.0012, 0.0015] and not resolved, in both cohorts (Figure 7); on PI error the interval included 0 although p was 0.047 (0.0024 [−0.0001, 0.0048]), and in the broad cohort it was resolved (0.0018 [0.0001, 0.0036]). The earlier operator gave a larger spatial nRMSE effect (0.0041 [0.0030, 0.0053]; Figure 7) than the block-centred operator; the corner-aligned minus block-centred difference was 0.0010 [0.0002, 0.0017], and the broad cohort agreed; on PI error the difference, 0.0016 [0.0003, 0.0028], had an interval excluding 0 with p = 0.062, and its interval included zero in the broad cohort.

**Figure 7.**
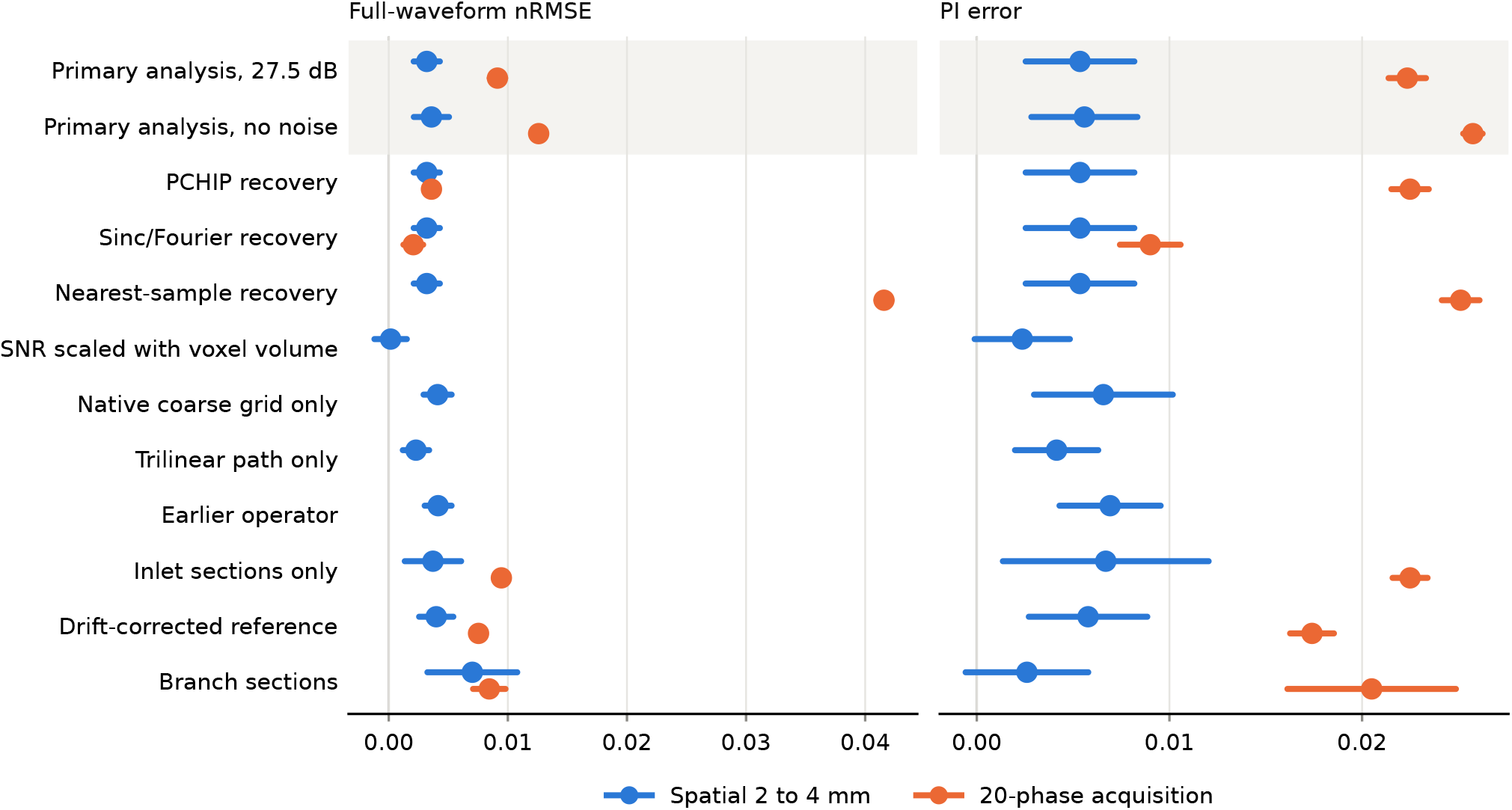
Spatial and 20-phase contrasts in the secondary and sensitivity analyses, primary cohort; the shaded row is the primary analysis. Points: family means; bars: t-based 95% CIs. The volume-scaled noise model, the single spatial paths and the earlier operator apply to the spatial contrast only.

The recovery method changed the 20-phase contrast but not the spatial one, which is evaluated at 80 frames (0.0032 for every method). The 20-phase nRMSE contrast was 0.0036 with PCHIP, 0.0021 with sinc/Fourier and 0.0416 with nearest-sample recovery, so the nRMSE ordering of the two contrasts depended on the method (Figure 7); no direct comparison was planned for the alternative methods. For PI error (Figure 7) and trough timing, the 20-phase point estimate exceeded the spatial one with every method in both cohorts. For peak timing it did so except with sinc/Fourier recovery, which lowered the peak timing error relative to the baseline condition (−0.25 [−0.37, −0.14] frames).

The remaining analyses shown in Figure 7 left both effects in the same direction (Supplementary Tables S12 and S14): the noise-free contrasts, each spatial path alone, the inlet sections only, the drift-corrected cycles, the broad cohort without the case with an inlet-ramp cycle, and the branch sections. The branch sections were available in 5 families, for which the sign-flip p value cannot fall below 0.062. Per-case cycle-closure values are listed in Supplementary Table S17.

### 3.5 Post hoc analyses

Two analyses were added post hoc and are descriptive. Relative to the baseline condition, sampling at 20 phases with point sampling, without averaging, increased nRMSE by 0.0065 [0.0059, 0.0071]; adding the box footprint at 20 phases increased it by a further 0.0026 [0.0025, 0.0028]. The signed PI change was negative under 20-phase acquisition in 7 of 7 families (mean −0.0273) and positive under spatial degradation in 6 of 7 (mean 0.0044); in the baseline condition, the signed PI error relative to the noise-free reference waveform ranged from 0.0007 to 0.0038.

## 4 Discussion

This benchmark measures how controlled degradations change the error of plane-flow endpoints computed from voxelised CFD fields, at the tested levels only. Within that scope, 20-phase acquisition changed PI error and diastolic-trough timing more than a 2 to 4mm spatial degradation, and the spatial effect on full-waveform nRMSE was small, consistent across geometries, and sensitive to the noise model and to the degradation operator.

The spatial degradation increased nRMSE by about 0.32% of the pulsatile range, in every family, and most of this was waveform-shape error: the change in cycle-mean flow was not resolved, and the cycle-mean flow bias carried 23% of the squared error. At 8mm this share rose to 60%: between 4 and 8mm, the cycle-mean flow error grew faster than the shape error. Phantom and CFD studies of the aorta and carotid bifurcation report that sensitivity to resolution depends on the quantity measured ^8–10^, and consensus statements ask for acquired and reconstructed resolution to be reported ^6,7^. The present data add that, for pulmonary-artery plane flow, doubling the voxel size from 2 to 4mm mainly distorts the waveform rather than the mean flow.

The principal comparisons were qualitatively similar when the broad cohort, which includes cases with numerical divergence, was analysed: every direct comparison had the same direction in both cohorts, and the broad-cohort intervals were in general wider.

The size of the spatial penalty depends on the noise model. With a fixed per-voxel SNR, the 4mm field carries the same relative noise as the 2mm field; when the SNR was scaled with voxel volume, as larger voxels would give at a fixed acquisition time, the nRMSE increase was no longer resolved. The benchmark does not model acquisition time, which in practice is traded against voxel size, phase count and averaging, so it cannot say which acquisition is preferable.

The effects of 20-phase acquisition on PI error and trough timing were robust: they exceeded the spatial effects with the primary recovery, and their point estimates remained larger with PCHIP, sinc/Fourier and nearest-sample recovery; for PI error this also held on the inlet sections alone, with drift correction and without noise. The trough and peak timing effects, about 8 and 9 ms with linear recovery, are fractions of the 50 ms sampling interval at 20 phases. The nRMSE ordering was not robust: it depended on the recovery method, and with sinc/Fourier recovery the 20-phase estimate was the smaller one in the primary cohort. Peak timing also depended on recovery. The 20-phase contrast combines fewer phases, one-interval averaging and the resulting noise reduction, in line with the distinction between nominal phase count and effective temporal averaging made in work on view sharing ^11–13^; the post hoc split attributed most of its nRMSE effect to the phase count and less to the averaging footprint.

The degradation operator itself changed the result. The earlier operator gave a larger spatial nRMSE effect than the block-centred one, although both apply the same point-spread function at the same nominal voxel size, so part of the effect measured with it reflects its sampling lattice. Studies of reconstruction and super-resolution methods report that image-level fidelity and derived haemodynamic quantities can diverge ^17–21^, and our previous study, which used the earlier operator, found that voxel fidelity did not rank PI accuracy ^14^; wherever such evaluations rely on synthetic degradation, the operator’s lattice is part of what they measure. Coarse samples should therefore sit at the centres of the regions they represent and temporal samples at their kernel centroids, and measurement planes should be placed by geometric criteria independent of the outcome, since planes that cross open CFD boundaries can create spurious features such as apparent flow reversal (Supplementary Note S6).

For studies that use degraded reference data to evaluate acquisitions or reconstruction methods, these results suggest what to report beyond the nominal voxel size and phase count: the baseline condition against which each contrast is measured, the spatial operator and its lattice, the effective temporal averaging window, the recovery method, and the noise model. Reporting the cycle-mean bias and the waveform-shape error separately would also show whether an nRMSE difference reflects a change in mean flow, which a plane-flow measurement would report directly, or a change in waveform shape, which affects PI and timing.

Three observations have possible mechanisms that this analysis did not test. Post hoc, the signed PI change was negative under 20-phase acquisition in 7 of 7 families and positive under spatial degradation in 6 of 7; one hypothesis for this post hoc observation is that box averaging and sampling at 20 phases attenuate the systolic peak and the diastolic trough, which lowers PI. The baseline condition contains noise, so its own errors are not zero and operators that average noise can offset part of their own error relative to it; this may explain why the 20-phase nRMSE contrast was larger without noise (0.0126) than at 27.5 dB (0.0091), and why the timing effects lie below the noise-free sampling-grid reference; neither explanation was tested. Finally, band-limited recovery can place an extremum between samples, and together with the noise averaging of the box footprint this may bring the recovered peak closer to the reference peak than in the baseline condition, which would give the negative peak-timing contrast seen with sinc/Fourier recovery.

## 5 Limitations

1. **CFD reference fields**. The simulations are not numerically converged (Supplementary Note S2), so the results describe how the operators degrade these fields, not errors against true haemodynamics. A verified CFD reference is planned.
2. **Non-periodic cycles**. The exported cycles are not periodic; the cycle-boundary step exceeded the 95th percentile of the in-cycle steps in 10 of 13 primary-cohort cases (Supplementary Table S17). The periodic interpolant spreads this step near the cycle boundary, where the trough lies; the drift correction bounds, but does not remove, this effect.
3. **Reference definition**. The reference is an unweighted bin average of cell-centre velocities, with filled core holes in 10 cases, and branch fluxes remain inflated relative to trunk flux at 2mm; branch sections are therefore secondary.
4. **Inflow**. The inlet boundary conditions give a cycle-mean inlet flow of 140–321 mL/s (8.4–19.3 L/min) in the primary cohort, higher than a typical resting pulmonary flow, and waveforms that stay well above zero in diastole (smallest trunk-section minimum, 68 mL/s; an example is shown in Figure 4). Because nRMSE is normalised by the pulsatile range and PI by the mean flow, both depend on the ratio of mean to pulsatile flow, which here differs from that of in vivo pulmonary flow; absolute error values and the bias share of the squared error may therefore not transfer to physiological waveforms.
5. **Precision**. With 7 families, effects smaller than about 1.27 between-family standard deviations cannot be expected to be detected with 80% power. Intervals are unadjusted for multiplicity, timing errors are integer-frame quantities, and part of the exclusion rule was written after the CFD-quality values were known.
6. **Plane coverage**. 66 of 88 candidate measurements met the geometric criteria, and branch sections exist only in some geometries, so the estimates are weighted towards the trunk and families contribute different mixes of sections.
7. **Integration**. Flow was integrated over the whole disk without a lumen mask, unlike region-of-interest flow quantification in clinical practice, which integrates within a segmented lumen contour; the noise in the disk margin adds to the error of every noisy condition.
8. **Point-spread function**. A single Gaussian point-spread function width was used, and other widths or shapes were not tested.
9. **Degradation and noise model**. The model is separable, with an isotropic point-spread function, temporal averaging of plane waveforms and no scanner forward model (k-space sampling, reconstruction, velocity encoding). Noise is Gaussian, independent per voxel and scaled with the signal rather than with a velocity-encoding setting, and the volume-scaled model ignores acquisition time. There are no in vivo or phantom data.

## 6 Conclusions

In this CFD benchmark, degrading the reference from 2 to 4mm increased full-waveform nRMSE by a small, consistent amount that was mostly waveform-shape error, and the increase was not resolved when the SNR scaled with voxel volume. Twenty-phase acquisition with one-interval averaging increased PI error and diastolic-trough timing error more than the spatial degradation, and its point estimates for these endpoints were larger with every recovery method; its nRMSE ordering and its effect on peak timing depended on the recovery method. The earlier operator gave a larger spatial effect than the block-centred one. These results describe benchmark behaviour on voxelised CFD fields, not clinical acquisition requirements.

## Data and code availability

The analysis code, the analysis plan, the case-exclusion rule and the aggregated results will be made publicly available on journal publication, and are available from the corresponding author on reasonable request. The geometries and CFD fields derive from patient data and are not distributed.

## Ethics statement

The SAPPHIRE study was conducted in accordance with the Declaration of Helsinki and approved by the Institutional Review Board of Hospital Universitario 12 de Octubre (protocol code 17/468). All patients gave written informed consent. The approval and consent covered the secondary research use of anonymised imaging data, including the derived computational geometries.

## Acknowledgments

This work was supported by ELKARTEK 2023 (CIC biomaGUNE and BCBL) and by the Basque Government, Department of Science, Universities and Innovation, through the IKUR 2030 Strategy, under the IKUR-HPC&AI 2025–2026 project SPADE-MRI (Synthetic Data Pipeline for Accelerated 4D Flow MRI: Enabling Robust Deep Learning Models). Jesús Ruiz-Cabello is funded by grant PID2024-155807OB-C21, funded by MCIN/AEI/10.13039/501100011033, by “ERDF A way of making Europe,” and by the European Union; grants DTS24/00043 and ISCIII-AES-2024/000225, funded by Instituto de Salud Carlos III (ISCIII); ELKARTEK 2025 (KK-2025-00084); and the Fundación contra la Hipertensión Pulmonar. CMR studies were acquired at the TRIMA@CNIC node of the ICTS ReDIB. This work acknowledges the use of ICTS-ReDIB, supported by the Ministry of Science, Innovation and Universities (MICIU) at BioImaC and CIC biomaGUNE. Computational resources were provided by the DIPC High-Performance Computing facility. We thank Juan F. Delgado (Hospital Universitario 12 de Octubre, Madrid) for access to the SAPPHIRE imaging data. We thank the DIPC HPC support team and collaborators at CIC biomaGUNE for technical discussions.

## Author contributions

Aitor Zubillaga-Unsain: Conceptualization, Methodology, Software, Investigation, Formal analysis, Data curation, Visualization, Writing–original draft, Writing–review & editing. Cesar Caballero-Gaudes: Funding acquisition, Resources, Supervision, Writing–review & editing. Jesús Ruiz-Cabello: Funding acquisition, Resources, Supervision, Writing–review & editing.

## Use of AI tools

AI assistants (OpenAI Codex/ChatGPT and Anthropic Claude, including Claude Code) were used extensively: to write and test the analysis code, run and document the analysis pipeline, draft the analysis plan, the case-exclusion rule and the manuscript text, check reproducibility, and carry out an AI-assisted critical review of an earlier draft, which identified interpretive issues and informed the design of the present analysis. All scientific decisions were made by the authors, who reviewed and verified the code, analyses and text, and take full responsibility for the content.

