## Supplementary Material for "Sensitivity of Pulmonary 4D Flow MRI Biomarkers to Controlled Spatial and Temporal Degradation in a CFD Benchmark"

Zubillaga-Unsain A, Caballero-Gaudes C, Ruiz-Cabello J. Preprint, version 1.

#### Note S1. Geometry families

The 22 cases comprise 11 geometries with two flow simulations each (Table S1). For 10 geometries the two flow simulations share a bit-identical source surface mesh; the remaining geometry was remeshed with an identical surface extent. The case-to-family mapping (hash `abb465a774d9`) is verified by every analysis script, and bootstrap resampling operates only on families. For analysis v2.5.0, Table S1 also lists each case's membership of the three cohorts of the exclusion rule, with the deciding criterion of each exclusion, and the number of geometric core holes filled in its 2 mm reference field.

**Table S1.** Geometry labels used in the main text, the corresponding case identifiers, cohort membership (analysis v2.5.0; one-pass application of the exclusion rule with amendment A1; the criterion in parentheses is the first failed criterion of that cohort) and the number of filled geometric core holes in the 2 mm reference field.

| Label | Case | Primary (non-divergent) | Broad | Fully verified | Filled holes |
|---|---|---|---|---|---|
| G1a | FFF | excluded (X0) | excluded (X0) | excluded (X0) | 0 |
| G1b | FFF-19 | retained | retained | excluded (B2) | 0 |
| G2a | FFF-1 | retained | retained | excluded (B2) | 0 |
| G2b | FFF-20 | retained | retained | excluded (B2) | 0 |
| G3a | FFF-2 | excluded (B′1) | retained | excluded (B′1) | 0 |
| G3b | FFF-21 | excluded (B′1) | retained | excluded (B′1) | 0 |
| G4a | FFF-3 | excluded (B′1) | retained | excluded (B′1) | 0 |
| G4b | FFF-22 | excluded (B′1) | retained | excluded (B′1) | 0 |
| G5a | FFF-4 | excluded (B′1) | retained | excluded (B′1) | 0 |
| G5b | FFF-23 | excluded (B′1) | retained | excluded (B′1) | 0 |
| G6a | FFF-5 | excluded (A) | excluded (A) | excluded (A) | 0 |
| G6b | FFF-24 | excluded (A) | excluded (A) | excluded (A) | 0 |
| G7a | FFF-6 | retained | retained | excluded (B2) | 12 |
| G7b | FFF-25 | retained | retained | excluded (B2) | 15 |
| G8a | FFF-7 | retained | retained | excluded (B2) | 3 |
| G8b | FFF-26 | retained | retained | excluded (B2) | 3 |
| G9a | FFF-8 | retained | retained | excluded (B2) | 32 |
| G9b | FFF-27 | retained | retained | excluded (B2) | 32 |
| G10a | FFF-9 | retained | retained | excluded (B2) | 6 |
| G10b | FFF-28 | retained | retained | excluded (B2) | 6 |
| G11a | FFF-10 | retained | retained | excluded (B2) | 1 |
| G11b | FFF-29 | retained | retained | excluded (B2) | 1 |

#### Note S2. Processing stages and methods details

Table S2 lists the processing stages of analysis v2.5.0; the details below complement the main-text Methods.

**Table S2.** Processing stages of analysis v2.5.0 and their resolution, taken from the frozen configuration and the data manifest.

| Stage | Resolution and definition |
|---|---|
| CFD solution | Fluent transient solutions on meshes of 0.76–0.98 million cells (equivalent cell edge 0.42–0.54 mm); second of two simulated cycles exported as 80 frames of 12.5 ms per 1.0 s cycle; time step 6.25 ms, except 12.5 ms in the run of case G1a (main text, Limitations) |
| Voxelised CFD reference field | 2 mm isotropic voxels; unweighted mean of the cell-centre velocities of the raw exported frames falling in each voxel bin (no scaling, no repair); vessel mask = voxels containing at least one cell centre; zero outside the mask |
| Core-hole filling | empty voxels whose centre lies inside the closed source surface and more than 16 mm from every open-cap vertex, filled by inverse-distance weighting (power 2) over the eight nearest raw cells and added to the mask; 111 voxels in 10 cases (Table S1); the fields of the other cases are identical to the unfilled binning |
| Degraded spatial levels | 4 mm (primary) and 8 mm (descriptive only) with the block-centred operator (Note S3); 4 mm with the earlier corner-aligned operator (sensitivity); each integrated on its native coarse grid or after trilinear interpolation to 2 mm |
| Noise | none; Gaussian, independent per voxel, component and frame, inside and outside the vessel mask, with standard deviation from the in-mask RMS velocity of the level; fixed per-voxel SNR of 27.5 and 17.5 dB (primary) or SNR scaled with voxel volume (45.56 and 35.56 dB at 4 mm, 63.62 and 53.62 dB at 8 mm); 10 realisations per noisy condition |
| Plane integration | disk aperture (maximal section radius + 8 mm); 96 radial × 192 angular midpoint quadrature; nearest-voxel lookup; no mask; the native coarse grid is passed with its exact origin and spacing |
| Temporal acquisition | 80 frames per cycle; 80, 40, 32, 25, 20, 16, 10 phases; centred point, box and triangle kernels on the band-limited periodic interpolant (unchanged from analysis v2.4.0) |

The reference is a property of the voxelised field. At 2 mm the summed branch flux of the reference field still exceeded its inlet flux in every case in which this was checked, although less than at 4 mm, and the voxelised trunk flux also differs from the flux computed directly from the raw cells; branch sections are therefore a secondary analysis. The 2 to 4 mm primary contrast uses the levels of our previous study [1].

*CFD runs.* All 22 runs used Ansys Fluent 2026 R1; the first flow simulation of each geometry (label a) used the SST $k$–$\omega$ turbulence model and the second (label b) was laminar, and the two were built from the same imported surface file. The inlet waveform is the same five-harmonic Fourier series in every run; it is scaled by 0.2 in every run except G1a, and in G3a it is also multiplied by a 2 s start-up ramp (Note S6). Walls are stationary and no-slip, with no mesh motion or fluid–structure coupling. Twenty of the 22 runs were driven by the same journal: two cycles with a time step of 6.25 ms, one inner iteration per time step, PISO pressure–velocity coupling with second-order pressure and first-order momentum, turbulence and time discretisation, and residual targets that were never met; their transcripts show no limiter messages. The other two runs were interactive and have no transcript: case G1a used a time step of 12.5 ms, up to 10 iterations per step and second-order time discretisation, without the inlet scale factor of the other cases, and case G2a used 6.25 ms, 15 iterations per step and second-order time discretisation. No case met the solver-convergence criterion of the exclusion rule (Note S7), so its fully verified cohort is empty. Every geometry has dome-shaped inlet caps, and inflow was prescribed as a velocity normal to the boundary. The solver-convergence test of the exclusion rule uses the continuity residuals recorded in the transcripts or residual monitors (Note S7).

*Illustrative case (main-text Figure 4).* The figure shows the inlet section of case G2a (selection and its timing in Note S8). The velocity maps show the noise-free through-plane velocity on the integration disk at frame 13, the systolic peak of the reference waveform, after block-centred degradation, on the native coarse grid of each level; no value in these maps is negative. They are sampled by nearest-voxel lookup, as in the plane integration, so voxel-layer boundaries appear as straight edges where the plane is oblique to the grid. The spatially degraded waveforms are at 80 frames and are the mean of the native-coarse-grid and trilinear paths; the 20-phase waveforms use box averaging at sampling offset 0, and their noise-free and 27.5 dB difference curves overlap. Noisy curves show the first of the 10 noise realisations. The interval between the last sample and frame 0 spans the cycle boundary (Table S17). The figure is descriptive and does not enter the inference.

*Integration.* Through-plane flow was integrated over each disk with a 96 radial × 192 angular polar midpoint quadrature, using nearest-voxel lookup without interpolation and no mask; in the reference field, voxels outside the vessel mask are zero, and after degradation the blurred velocity and the noise within the whole disk, including its margin outside the lumen, are integrated. On the native coarse grid the integrator receives origin $(f-1)h/2$ and spacing $fh$.

*Noise realisations.* Each noise field has its own seed, derived from a cryptographic hash of a base seed (20260622), the case, the spatial level, the noise model, the noise level and the realisation; every seed is recorded with the results. Within a spatial level, noise model, noise level and realisation, one noise field is shared by all temporal conditions, spatial paths and recovery methods, and different levels have independent noise. The case value of a noisy condition is the mean over its 10 realisations; noise-free conditions have one.

*Temporal sampling.* For $T$ phases per cycle and offset fraction $r$ (0, 0.25, 0.5 or 0.75 of the output interval), sample $k$ has the nominal time $c_k = (k+r)N/T$ ($N = 80$ frames) and is the kernel-weighted average of the periodic band-limited interpolant of the 80-frame plane waveform, centred on $c_k$. Estimates for $T < 80$ average over the four offsets.

*Spatial paths.* The native-grid and trilinear paths were weighted equally in every estimate; at 2 mm both paths are the identity.

*Offsets and timing.* At 80 frames, estimates use offset 0 only, the only 80-frame condition in which the temporal operator is an exact identity; non-zero offsets at 80 frames enter operator checks only. Peak and trough times are the frames of the maximum and minimum of the waveform, with ties resolved by taking the first frame, and the timing error is the unsigned periodic distance between recovered and reference times.

*Drift correction.* The linear drift correction of the sensitivity analysis is $q'(k) = q(k) + \frac{k+1}{80}D$ with $D = \frac{80}{79}\big(q(0) - q(79) - s\big)$, where $s$ is the mean of the first and last in-cycle steps. It was applied to each 80-frame plane waveform, the reference and every degraded waveform alike, before temporal acquisition, using that waveform's own first and last frames, with periodic-linear recovery only, and it was computed as a separate grid.

*Endpoint constants and statistics.* Full-waveform nRMSE is $\mathrm{RMSE}(Q_{\mathrm{pred}} - Q_{\mathrm{ref}})/\max(P_{95} - P_5, \delta_Q)$ and PI is $(Q_{\mathrm{sys}} - Q_{\mathrm{dia}})/\max(|Q_{\mathrm{mean}}|, \delta_{PI})$. The denominator floors $\delta_Q$ and $\delta_{PI}$ (Table S3) were derived from the noise-free reference waveforms of all 22 cases before any degraded output existed. Case G1a, although excluded from every cohort, contributes to them (about 16% of the amplitude scale); no floor is active with or without it. The family bootstrap uses 10,000 resamples of families with replacement (seed 20260625) and percentile intervals. The descriptive endpoints (unsigned systolic, diastolic and mean-flow errors, and the waveform correlation) are available with the analysis outputs.

**Table S3.** Endpoint denominator floors, derived from the noise-free reference waveforms before any degraded output was computed. Neither floor was active on any reference plane.

| Constant | Value | Construction | Active on any reference plane? |
|---|---|---|---|
| $\delta_Q$ (nRMSE denominator floor) | 2.114 mL/s | 1% of the median reference P95–P5 range | no (smallest reference range (max − min) 67.9 mL/s) |
| $\delta_{PI}$ (PI denominator floor) | 2.114 mL/s | 1% of the same scale | no (smallest reference $|Q_{\mathrm{mean}}|$ 79.8 mL/s) |

### Note S3. Spatial operator and its verification

The spatial operator of analysis v2.4.0, also used in our previous study [1], filtered the field with a Gaussian point-spread function and then downsampled it by trilinear interpolation with corner alignment. A registration check of the degraded voxels, run before any degraded output of analysis v2.5.0 existed, showed that the integrator coordinates equal the positions of that operator's samples, so its degraded field is not misregistered against the planes. Its lattice, however, is anchored at the first and last reference voxel of each axis: its pitch exceeds the nominal voxel size, differently on each axis, and its samples are displaced by up to about one coarse voxel from a block-centred acquisition grid, with a degraded mask that is offset from the velocity samples. The operator was therefore replaced before any degraded output existed, and kept as a sensitivity condition at 4 mm.

In the block-centred operator, for a coarsening factor $f$, coarse voxel $j$ covers reference voxels $[fj, f(j+1))$ along each axis, with centre $c_j = fj + (f-1)/2$ in reference index units. Its value is the reference field filtered by separable Gaussian weights $w(m) \propto \exp(-(m-c_j)^2/2\sigma^2)$ for $|m - c_j| \leq 4\sigma$, with $\sigma = 0.4f$ reference voxels, normalised over the full truncated support, and sampled at the block centre without block averaging, so that the effective spatial response is exactly the pre-declared Gaussian. Incomplete trailing blocks are zero-padded (the true exterior value), giving $\lceil N/f \rceil$ coarse voxels per axis; at 4 mm no vessel voxel falls in an incomplete block, at 8 mm some do. A coarse voxel is in the degraded mask if any reference voxel of its padded block is in the reference mask; the mask sets the noise level and is not used by the integrator. On the native path the integrator receives origin $(f-1)h/2$ and spacing $fh$; the trilinear path interpolates separably from the coarse centres to the reference voxel centres, with edge replication outside the range of the coarse centres.

The operators were verified by the gates in Table S4. Gates A1–A6 use synthetic fields only and passed (commit `7b19415194c5`). Gates B1–B5 were run on six pilot cases at every spatial level, and on the corresponding pilot of the drift-corrected reference; they read the pilot outputs for operator checks only (pass or fail and operator deviations), not for any endpoint summary. B1–B4 passed in every case. B5 failed on both pilots in its sub-check that the unsigned mean-flow error equals the absolute signed error; after the failure and a diagnostic limited to failure counts and deviation magnitudes had been seen, and before any endpoint summary, the tolerance of that comparison was changed to a combined tolerance relative to the reference mean flow and to the signed error (author decision; commit `84f322ff1863`). With it, every gate passed on both pilots, and the full grids were then run. No gate was run on the full grids (Note S8).

**Table S4.** Verification gates of the spatial operators and the grid (analysis v2.5.0).

| Gate | Check | Outcome |
|---|---|---|
| A1 | coarse spacing exactly 4 and 8 mm on every axis, for every case shape | pass (synthetic) |
| A2 | block centres equal the integrator coordinates | pass (synthetic) |
| A3 | uniform and linear fields reproduced on the native and trilinear paths | pass (synthetic) |
| A4 | flux of a uniform field through a synthetic plane equals $U\pi R^2$ | pass (synthetic) |
| A5 | impulse response equals the pre-declared normalised Gaussian at the block centre | pass (synthetic) |
| A6 | earlier corner-aligned operator bit-identical to the code of analysis v2.4.0 | pass (synthetic) |
| B1 | identity: at 2 mm without noise, 80 frames, point sampling and offset 0, the nRMSE, mean-removed nRMSE and signed mean-flow error vanish to numerical precision and the timing errors are zero | pass (pilots) |
| B2 | temporal attenuation against an independent quadrature of the band-limited interpolant (sinc and $\mathrm{sinc}^2$ gains) | pass (pilots) |
| B3 | integrator origin and spacing per level, planes, and reference cycle-mean flows equal to the constants file | pass (pilots) |
| B4 | provenance: hashes, commit, operator versions, recomputed noise seeds, row counts | pass (pilots) |
| B5 | endpoint computation: exact bias/shape identity, unsigned equal to absolute signed mean-flow error, bias share in $[0, 1]$ | pass (pilots) after the tolerance change |

### Note S4. Geometric plane placement

Planes were placed on the exact source surfaces (wall and labelled open caps) by cross-section tracking from each cap, with criteria P1–P8 (main text, Measurement planes). The rule is that of analysis v2.4.0 (hash `aef087140c43`), used unchanged except for one amendment of P8. It was written and placed under version control before it was applied to any case of analysis v2.4.0 and before any flow was computed on the resulting planes. Three implementation corrections were made then, before any flow computation, with no threshold changed:

1. an indexing fix for section chains clipped by the search radius;
2. robust entry from curved caps, and suppression of the bifurcation stop while the section still contained cap segments;
3. locating the bifurcation from the first meeting of branch and trunk tracks.

Each correction was triggered by geometric quality control and is recorded with its reason in the version-controlled rule file.

In analysis v2.5.0 only two steps depend on the grid: P8 is evaluated on the 2 mm reference mask (the binning mask with the filled holes), and plane centres are converted to the integrator frame with $h = 2$ mm. A first run with the unamended rule lost nine measurements that were valid in v2.4.0; each failed P8 only because of a small sampled mask region lying outside the wall, with no sample point inside the closed source surface. The amendment (hash `ab95c722f56f`) therefore ignores, when P8 counts regions, any sampled mask region none of whose points lies inside the closed source surface; the test on the outer ring of the disk still uses every sampled mask point, and nothing else changed. It was decided after the geometric P8 diagnostic of the unamended set and the reference waveforms of that set, used for a first and superseded version of the endpoint constants, had been seen, and before any degraded output existed. With it, every candidate measurement with a geometrically valid station is included (66 of 88), at the same station as in analysis v2.4.0 to cross-platform rounding, and the resulting plane set was fixed before any degraded output was computed; four measurements that failed P8 on the 4 mm grid of v2.4.0 are gained (the mid-trunk section of G1 and the first outlet of G6, in both flow simulations). Table S5 lists validity by geometry, which is identical for both flow simulations of a geometry. The frozen plane set has hash `5d69d05db17f`.

**Table S5.** Measurement validity by geometry, analysis v2.5.0 (66 of 88 candidate measurements).

| Geometry | Inlet | Mid | Outlet 1 | Outlet 2 | Exclusion reason |
|---|---|---|---|---|---|
| G1 | ✓ | ✓ | ✓ | ✓ | – |
| G2 | ✓ | ✓ | ✓ | ✓ | – |
| G3 | ✓ | ✓ | – | – | outlet 1: no geometrically valid section; outlet 2: no geometrically valid section |
| G4 | ✓ | ✓ | – | – | outlet 1: no geometrically valid section; outlet 2: no geometrically valid section |
| G5 | ✓ | ✓ | ✓ | ✓ | – |
| G6 | ✓ | ✓ | ✓ | ✓ | – |
| G7 | ✓ | ✓ | – | – | outlet 1: no geometrically valid section; outlet 2: no geometrically valid section |
| G8 | ✓ | – | ✓ | ✓ | mid: no geometrically valid section |
| G9 | ✓ | ✓ | ✓ | – | outlet 2: no geometrically valid section |
| G10 | ✓ | – | – | – | mid: no geometrically valid section; outlet 1: no geometrically valid section; outlet 2: no geometrically valid section |
| G11 | ✓ | ✓ | ✓ | ✓ | – |

### Note S5. Temporal operator verification

The temporal operator is unchanged from analysis v2.4.0, and its tests were re-run in the pilot gate jobs of this version. They use analytic periodic signals (sinusoid, off-grid-shifted sinusoid, four-harmonic waveform) for every phase count, footprint and offset, and check:

- kernel centroid equal to the nominal sample time;
- unit kernel mass;
- exact requested width;
- zero phase delay with sinc recovery;
- agreement with an independent brute-force quadrature, including periodic wrap-around;
- exact identity at 80 frames, point sampling and offset 0.

Periodic-linear recovery carries only a bounded second-order phase term of the interpolator itself, of at most 0.003 frames. A Nyquist component sampled at a fractional offset cannot be recovered uniquely, which affects sinc recovery at 80 frames with non-zero offsets; these conditions enter operator checks only. Pilot gates B1 and B2 (Table S4) check the identity and the temporal attenuation on the pilot cases.

### Note S6. Reference-data quality

- *Cycle closure.* No case is cycle-converged after two simulated cycles. The cycle-closure metric, the step between the last and first frame of the reference waveform divided by the 95th percentile of its in-cycle steps and averaged over the trunk sections, exceeded 1 in 10 of 13 primary-cohort cases (range 0.75–1.51) and in 15 of 19 broad-cohort cases (range 0.75–5.10; Table S17). The linear drift correction of the sensitivity analysis is defined in the main text.
- *Inlet-cap runaway.* In eight cases of four geometries (G3–G6), cells next to the inlet cap reached speeds above the ceiling of criterion $B'1$ (Table S7); these cases are excluded from the primary cohort. Earlier products of the dataset replaced some of these samples by temporal interpolation; no repaired or rescaled field is used in this analysis.
- *Case G3a.* Its exported cycle lies under a documented inlet ramp of 2 s (criterion F), so its source waveform is not a periodic cycle and differs from that of the second flow simulation of the geometry. G3a fails $B'1$, $B'2$ and F and is excluded from the primary cohort; it is retained in the broad cohort, and the broad-cohort analysis without it is reported in the main text.

- *Case G1a.* Its raw export corresponds to an inflow five times that of the other cases, and it is excluded from every cohort (criterion X0). Its unscaled reference waveforms contribute to the endpoint constants; no denominator floor is active with or without them (main text).
- *Flow direction.* On every reference plane the reference waveform is forward throughout the cycle (largest reverse-flow fraction 0.000, against a threshold of 0.02 for substantial reverse flow), and every cycle-mean flow is positive. An apparent flow reversal seen at two inlets in an earlier, non-geometric plane placement was caused by integration disks crossing the open inlet cap; it is not a property of the flow.

### Note S7. Case-exclusion rule and amendment A1

The exclusion rule (hash `ea5f3d46e7d7`) uses only the quality of the CFD solution as exported: the raw, unscaled and unrepaired cell-centre frames. It never uses benchmark endpoints, degraded data or any quantity computed by the benchmark pipeline, and it is applied by one script, in one pass, to all 22 cases. Table S6 lists its criteria, their thresholds and the cohorts in which each applies (amendment A1, hash `66a4dc75c73a`). A criterion that cannot be evaluated for a case neither passes nor fails it, and a family is retained if at least one of its cases is retained. Table S7 gives the per-case outcomes; the resulting cohort membership is in Table S1.

**Table S6.** Criteria of the case-exclusion rule, their thresholds, and the cohorts in which they apply (amendment A1).

| Criterion | Definition | Pass if | Primary | Broad | Fully verified |
|---|---|---|---|---|---|
| X0 | explicit exclusion of case G1a (raw export at five times the inflow of the other cases) | – | ✓ | ✓ | ✓ |
| A | maximum speed outside the cap zones (cells within 16 mm of an open-cap vertex) | ≤ 3 m/s | ✓ | ✓ | ✓ |
| B′1 | maximum speed of any cell in the domain, in any exported frame | ≤ 10 m/s | ✓ | – | ✓ |
| B′2 | sustained growth of the domain maximum speed over the last 10 frames to at least twice its median | absent | ✓ | – | ✓ |
| B2 | end-of-step continuity residuals, where solver logs exist | $\leq 10^{-2}$ | – | – | ✓ |
| C | inlet vs mid-trunk cycle-mean flow (not evaluable without a mid-trunk section) | ≤ 5% | ✓ | ✓ | ✓ |
| D1 | maximum speed on the inlet section | ≤ 3 m/s | ✓ | ✓ | ✓ |
| D2 | phase-resolved inlet vs mid-trunk flow disagreement, relative to the pulsatile range (not evaluable without a mid-trunk section) | ≤ 10% | ✓ | ✓ | ✓ |
| E1 | criteria judged on the raw, unrepaired, unscaled solution; repair cannot rescue a case | – | ✓ | ✓ | ✓ |
| E2 | cap-zone cells above the speed ceiling of A lie far from every section | > 10 mm | ✓ | ✓ | ✓ |
| F | exported cycle overlaps an inlet ramp or start-up documented in the case setup (not evaluable without setup documentation) | no overlap | ✓ | – | ✓ |

**Table S7.** Per-case outcomes of the one-pass rule application: maximum speed outside the cap zones (A) and in the whole domain (B′1), and the outcome of every other criterion. Bold: failed; n.e.: not evaluable.

| Label | Case | A (m/s) | B′1 (m/s) | B′2 | B2 | C | D1 | D2 | E2 | F | X0 |
|---|---|---|---|---|---|---|---|---|---|---|---|
| G1a | FFF | **5.88** | 7.74 | pass | **fail** | n.e. | **fail** | n.e. | **fail** | n.e. | **fail** |
| G1b | FFF-19 | 1.11 | 1.63 | pass | **fail** | n.e. | pass | n.e. | pass | pass | – |
| G2a | FFF-1 | 0.86 | 1.07 | pass | **fail** | pass | pass | pass | pass | n.e. | – |
| G2b | FFF-20 | 0.76 | 1.22 | pass | **fail** | pass | pass | pass | pass | pass | – |
| G3a | FFF-2 | 0.55 | **38** | **fail** | **fail** | pass | pass | pass | pass | **fail** | – |
| G3b | FFF-21 | 0.81 | **676** | pass | **fail** | pass | pass | pass | pass | pass | – |
| G4a | FFF-3 | 0.83 | **31** | pass | **fail** | pass | pass | pass | pass | pass | – |
| G4b | FFF-22 | 0.83 | **54** | pass | **fail** | pass | pass | pass | pass | pass | – |
| G5a | FFF-4 | 2.21 | **236** | pass | **fail** | pass | pass | pass | pass | pass | – |
| G5b | FFF-23 | 2.22 | **296** | pass | **fail** | pass | pass | pass | pass | pass | – |
| G6a | FFF-5 | **3.26** | **2163** | pass | **fail** | pass | pass | pass | **fail** | pass | – |
| G6b | FFF-24 | **3.42** | **2201** | pass | **fail** | pass | **fail** | pass | **fail** | pass | – |
| G7a | FFF-6 | 1.10 | 3.71 | pass | **fail** | pass | pass | pass | pass | pass | – |
| G7b | FFF-25 | 0.94 | 3.03 | pass | **fail** | pass | pass | pass | pass | pass | – |
| G8a | FFF-7 | 0.79 | 0.81 | pass | **fail** | n.e. | pass | n.e. | pass | pass | – |
| G8b | FFF-26 | 0.79 | 0.81 | pass | **fail** | n.e. | pass | n.e. | pass | pass | – |
| G9a | FFF-8 | 1.20 | 2.00 | pass | **fail** | pass | pass | pass | pass | pass | – |
| G9b | FFF-27 | 1.20 | 2.00 | pass | **fail** | pass | pass | pass | pass | pass | – |
| G10a | FFF-9 | 0.72 | 1.03 | pass | **fail** | n.e. | pass | n.e. | pass | pass | – |
| G10b | FFF-28 | 0.72 | 1.03 | pass | **fail** | n.e. | pass | n.e. | pass | pass | – |
| G11a | FFF-10 | 0.61 | 0.93 | pass | **fail** | pass | pass | pass | pass | pass | – |
| G11b | FFF-29 | 0.61 | 0.93 | pass | **fail** | pass | pass | pass | pass | pass | – |

The record of the rule is as follows. The thresholds of A, B2, C, D and E were written in a first draft before any case value existed and were not changed. Per-case values of A, D and E, and of a first divergence criterion (B1), were then computed and seen. B1 was withdrawn because it measured cycle closure rather than runaway growth, and B′ and F, including their thresholds, were written after those values had been seen, before B′ and F themselves were computed; a provenance check proposed for F was later withdrawn as an exclusion test for the same reason (construct validity). Three clarifications made after the values were seen define metrics and the handling of cases, not thresholds. Amendment A1 was decided after these facts, the solver-residual facts and a provenance audit of the runs were known, and before any pilot or grid result was read. It changed the evidence for F to the setup documentation recorded in the audit, because the intended provenance table has no ramp or exported-window field (the links from each exported frame set to its run are inferred in the audit, not hashed), and it replaced the rule's two cohorts by the three used here. When the rule was applied, B2 was judged failed where any documented continuity residual of the run exceeds $10^{-2}$; the growth ratio of B2 cannot be computed from the four-step residual excerpts available and is not needed for a failure. This reading was recorded before unsealing and affects only the fully verified cohort, which is empty either way. An erratum records that the criterion values were computed one day earlier than the frozen documents state; no value, outcome or cohort changes.

### Note S8. Provenance and analysis history

*Analysis history.* The first internal analysis of the same cases (v2.3.1) used a temporal operator with sample-timing bias (sample times rounded down to whole frames, and even-width averaging windows not centred on the sample time) and plane acceptance tuned on outcomes. It was superseded before the plan of the next analysis was written, and none of its results are reported. That next analysis, v2.4.0, used a 4 mm reference field with a 4 to 8 mm spatial contrast, one noise realisation per condition, all 22 cases and family-bootstrap intervals. It was the basis of an earlier, unpublished draft of this manuscript, which underwent an AI-assisted critical review; the review identified interpretive issues, led to post hoc analyses of that version, and informed the design of the present analysis. Analysis v2.4.0 is superseded and none of its results are reported. A spatial–temporal interaction analysis pre-declared in the v2.4.0 plan was withdrawn after review because its log-offset constant is not dimensionally consistent across endpoints; it is not part of the present plan. The present design was therefore not blind to either earlier analysis.

*Identifiers of analysis v2.5.0* (hash prefixes):

- analysis plan `567f06eb3b34` (commit `0b2473b89d7c`); exclusion rule `ea5f3d46e7d7`; amendment A1 `66a4dc75c73a` (commit `339a58088069`); one-pass rule application commit `2f576e6e21c3`;

- grid code commit `ead88cc8d354`; grid configuration `b811d46c0dd6`; drift-corrected grid code commit `37079863add3`;
- constants `be5e52883753`; plane set `5d69d05db17f`; plane-rule amendment `ab95c722f56f`; data manifest `9a046796f57b`; family map `1cb94d056374`;
- analysis script commit `ff0d1c55d55c` (script `904cfdce4998`); post hoc signed-PI script `3e6f5af16e50`;
- noise seed base 20260622 and bootstrap seed 20260625.

*Sealing and unsealing record.* An append-only sealing log records the frozen artefacts, every grid start and end time and every access to a results file; copies at three dates are in the repository: `results/paper2_v2_5_0/SEALING_LOG.md` (hash `c10963523029`), `results/paper2_v2_5_0/SEALING_LOG_update1.md` (hash `bf78fb96779a`), which adds the entries for the illustrative recomputation below, and `results/paper2_v2_5_0/SEALING_LOG_update2.md` (hash `2b57d705493f`), which adds a correction of one of those entries. In order:

1. The analysis plan, the exclusion rule with amendment A1, and the computational configuration were frozen under version control and pushed before any pilot or grid result was read.
2. The one-pass application of the exclusion rule was committed and pushed while no results file had yet been read.
3. The analysis script was written against the plan and tested on synthetic rows only (commit `9affc5bb1143`).
4. The pilot outputs were read by the operator gates only; after the B5 tolerance change (Note S3) every gate passed, and the main grid and the drift-corrected grid were then run and sealed.
5. Before unsealing, it was recorded (author decision) that no gate would be run on the full grids: the only gates defined are the pilot gates, and the plan's condition that the drift-corrected grid has passed its gates was read as referring to them.
6. The analysis script was then extended with a startup verification of the frozen inputs (plan, rule, amendment and cohort table) and a structural pre-check of the grid outputs, without any change to statistical or endpoint code; its tests passed in the environment of the analysis job, and unsealing was then approved (author decision).
7. The analysis job logged every results-file read. The unit tests of the grid and operator code were re-run on the analysis commit after unsealing, because they had last run at the gate-fix commit; a test file that the gate jobs had omitted from their test call was added for later runs (commit `f846889d63d0`), without affecting any run already made.
8. The post hoc signed-PI analysis was run after unsealing by a separate script whose reads were logged.
9. After unsealing, the waveforms of one case (G2a, inlet section) were recomputed with the code and configuration of the main grid for the illustrative main-text figure (Figure 4; details in Note S2); the recomputation reads no results file, and no number in the text depends on it. The case was selected after the aggregate results had been unsealed and before any waveform of it was recomputed or plotted; the log entry that starts the recomputation names it. It is the first flow simulation of the first geometry whose two flow simulations are both in the primary cohort; this description of the choice was recorded later, with the addition of the figure, and is not a rule stated before it. An earlier log statement that the case was chosen before any waveform was computed is corrected in the log: the main grid computes every plane waveform internally, although it stores only endpoint values.
10. An erratum corrects the recorded date of the CFD-quality values (commit `5675c65d611b`; Note S7).

Every number, table and figure in the manuscript and this supplement is generated from the saved results and the frozen configuration by versioned scripts.

#### Note S9. Analysis v2.5.0: full results

Tables S8–S14 give the estimates of the main-text figures for both cohorts. Estimates are family means of paired differences with t-based 95% CIs and exact sign-flip p values; bold: CI excludes 0; ‡: the CI and the p value disagree (26 of 566 tested summaries); §: descriptive only. Primary: non-divergent cohort (7 families); broad: 10 families. CIs and p values are unadjusted for multiplicity.

**Table S8.** Pre-specified contrasts: full-waveform nRMSE and PI error.

| Contrast | nRMSE: primary | nRMSE: broad | PI: primary | PI: broad |
|---|---|---|---|---|
| Spatial (4 vs 2 mm) | **0.0032 [0.0021, 0.0043]** $p = 0.016$ | **0.0032 [0.0023, 0.0042]** $p = 0.002$ | **0.0054 [0.0025, 0.0082]** $p = 0.016$ | **0.0044 [0.0023, 0.0065]** $p = 0.002$ |
| Temporal acquisition (20 phases, box) | **0.0091 [0.0085, 0.0098]** $p = 0.016$ | **0.0110 [0.0068, 0.0153]** $p = 0.002$ | **0.0223 [0.0214, 0.0233]** $p = 0.016$ | **0.0298 [0.0161, 0.0436]** $p = 0.002$ |
| Footprint (triangle vs box) | **0.0033 [0.0032, 0.0033]** $p = 0.016$ | **0.0032 [0.0031, 0.0033]** $p = 0.002$ | **0.0130 [0.0123, 0.0136]** $p = 0.016$ | **0.0138 [0.0123, 0.0152]** $p = 0.002$ |
| Noise (17.5 vs 27.5 dB) | **0.0078 [0.0066, 0.0090]** $p = 0.016$ | **0.0083 [0.0069, 0.0098]** $p = 0.002$ | **0.0114 [0.0075, 0.0154]** $p = 0.016$ | **0.0117 [0.0088, 0.0146]** $p = 0.002$ |
| Combined (4 mm, 20 phases, box) | **0.0095 [0.0086, 0.0104]** $p = 0.016$ | **0.0118 [0.0071, 0.0164]** $p = 0.002$ | **0.0200 [0.0153, 0.0246]** $p = 0.016$ | **0.0294 [0.0137, 0.0451]** $p = 0.002$ |
| Spatial (8 vs 2 mm)[§] | 0.0094 descriptive | 0.0104 descriptive | 0.0075 descriptive | 0.0076 descriptive |

**Table S9.** Pre-specified contrasts: diastolic-trough and systolic-peak timing error (frames of 12.5 ms).

| Contrast | Trough: primary | Trough: broad | Peak: primary | Peak: broad |
|---|---|---|---|---|
| Spatial (4 vs 2 mm) | 0.04 [−0.04, 0.11] $p = 0.344$ | 0.03 [−0.02, 0.09] $p = 0.203$ | 0.14 [−0.01, 0.29] $p = 0.094$ | **0.15 [0.05, 0.26]** $p = 0.016$ |
| Temporal acquisition (20 phases, box) | **0.63 [0.52, 0.74]** $p = 0.016$ | **0.76 [0.56, 0.97]** $p = 0.002$ | **0.72 [0.65, 0.79]** $p = 0.016$ | **0.72 [0.67, 0.78]** $p = 0.002$ |
| Footprint (triangle vs box) | 0.04 [−0.03, 0.11] $p = 0.125$ | 0.03 [−0.01, 0.07] $p = 0.062$ | 0.000 [0.000, 0.000] $p = 1.000$ ¶ | 0.002 [−0.003, 0.008] $p = 1.000$ ¶ |
| Noise (17.5 vs 27.5 dB) | **0.23 [0.16, 0.30]** $p = 0.016$ | **0.24 [0.19, 0.29]** $p = 0.002$ | **0.30 [0.18, 0.43]** $p = 0.016$ | **0.35 [0.25, 0.45]** $p = 0.002$ |
| Combined (4 mm, 20 phases, box) | **0.63 [0.52, 0.74]** $p = 0.016$ | **0.77 [0.55, 0.99]** $p = 0.002$ | **0.72 [0.65, 0.79]** $p = 0.016$ | **0.72 [0.67, 0.78]** $p = 0.002$ |
| Spatial (8 vs 2 mm)[§] | 0.13 descriptive | 0.12 descriptive | 0.21 descriptive | 0.23 descriptive |

[¶]Printed with 3 decimals (all other values: 2) because the effect is below 0.005 frames in both cohorts.

**Table S10.** Direct comparisons between the spatial and 20-phase contrasts (paired differences within family; periodic-linear recovery).

| Comparison (periodic-linear recovery) | Primary cohort | Broad cohort |
|---|---|---|
| nRMSE: spatial − temporal acquisition | **−0.0059 [−0.0074, −0.0045]** $p = 0.016$ | **−0.0078 [−0.0117, −0.0040]** $p = 0.002$ |
| PI: spatial − temporal acquisition | **−0.0170 [−0.0199, −0.0141]** $p = 0.016$ | **−0.0254 [−0.0399, −0.0110]** $p = 0.002$ |
| Trough timing: temporal acquisition − spatial | **0.59 [0.48, 0.70]** $p = 0.016$ | **0.73 [0.52, 0.94]** $p = 0.002$ |
| Peak timing: temporal acquisition − spatial | **0.58 [0.49, 0.67]** $p = 0.016$ | **0.57 [0.50, 0.65]** $p = 0.002$ |
| Mean-removed nRMSE: spatial − temporal acquisition | **−0.0066 [−0.0078, −0.0055]** $p = 0.016$ | **−0.0087 [−0.0129, −0.0045]** $p = 0.002$ |

**Table S11.** Bias/shape decomposition: signed change in cycle-mean flow (% of the reference), mean-removed nRMSE and the bias share of the squared error of the test condition.

| Contrast | Signed mean flow (%): primary | broad | Mean-removed nRMSE: primary | broad | Bias share (%): primary | broad |
|---|---|---|---|---|---|---|
| Spatial (4 vs 2 mm) | 0.08 [−0.09, 0.24]<br>$p = 0.328$ | 0.01 [−0.13, 0.14]<br>$p = 0.938$ | **0.0025 [0.0017, 0.0033]**<br>$p = 0.016$ | **0.0023 [0.0017, 0.0030]**<br>$p = 0.002$ | 23 | 29 |
| Temporal acquisition (20 phases, box) | 0.00 [0.00, 0.00]<br>$p = 1.000$ | 0.00 [0.00, 0.00]<br>$p = 1.000$ | **0.0091 [0.0085, 0.0098]**<br>$p = 0.016$ | **0.0111 [0.0068, 0.0154]**<br>$p = 0.002$ | 0 | 0 |
| Footprint (triangle vs box) | 0.00 [0.00, 0.00]<br>$p = 1.000$ | 0.00 [0.00, 0.00]<br>$p = 1.000$ | **0.0033 [0.0032, 0.0033]**<br>$p = 0.016$ | **0.0032 [0.0031, 0.0033]**<br>$p = 0.002$ | 0 | 0 |
| Noise (17.5 vs 27.5 dB) | −0.01 [−0.02, 0.01]<br>$p = 0.297$ | 0.00 [−0.02, 0.01]<br>$p = 0.424$ | **0.0078 [0.0066, 0.0089]**<br>$p = 0.016$ | **0.0083 [0.0069, 0.0097]**<br>$p = 0.002$ | 1 | 1 |
| Combined (4 mm, 20 phases, box) | 0.08 [−0.09, 0.24]<br>$p = 0.328$ | 0.01 [−0.13, 0.14]<br>$p = 0.938$ | **0.0090 [0.0080, 0.0101]**<br>$p = 0.016$ | **0.0113 [0.0067, 0.0159]**<br>$p = 0.002$ | 7 | 5 |
| Spatial (8 vs 2 mm)§ | −0.04<br>descriptive | −0.34<br>descriptive | 0.0056<br>descriptive | 0.0059<br>descriptive | 60 | 61 |

**Table S12.** Secondary analyses: branch sections, noise-free contrasts, single spatial paths and the volume-scaled noise model.

| Secondary analysis | nRMSE: primary | nRMSE: broad | PI: primary | PI: broad |
|---|---|---|---|---|
| Branch sections: spatial | **0.0070 [0.0032, 0.0108]**<br>$p = 0.062^{\ddagger}$ | **0.0072 [0.0043, 0.0101]**<br>$p = 0.031$ | 0.0026 [−0.0006, 0.0058]<br>$p = 0.125$ | **0.0032 [0.0003, 0.0061]**<br>$p = 0.062^{\ddagger}$ |
| Branch sections: temporal acquisition | **0.0084 [0.0071, 0.0098]**<br>$p = 0.062^{\ddagger}$ | **0.0085 [0.0074, 0.0095]**<br>$p = 0.031$ | **0.0205 [0.0161, 0.0249]**<br>$p = 0.062^{\ddagger}$ | **0.0217 [0.0171, 0.0264]**<br>$p = 0.031$ |
| Noise-free: spatial | **0.0036 [0.0021, 0.0051]**<br>$p = 0.016$ | **0.0040 [0.0026, 0.0054]**<br>$p = 0.002$ | **0.0056 [0.0028, 0.0083]**<br>$p = 0.016$ | **0.0054 [0.0036, 0.0072]**<br>$p = 0.002$ |
| Noise-free: temporal acquisition | **0.0126 [0.0123, 0.0128]**<br>$p = 0.016$ | **0.0148 [0.0101, 0.0194]**<br>$p = 0.002$ | **0.0258 [0.0252, 0.0263]**<br>$p = 0.016$ | **0.0336 [0.0192, 0.0480]**<br>$p = 0.002$ |
| Native path only: spatial | **0.0041 [0.0029, 0.0053]**<br>$p = 0.016$ | **0.0041 [0.0032, 0.0050]**<br>$p = 0.002$ | **0.0066 [0.0030, 0.0102]**<br>$p = 0.016$ | **0.0056 [0.0030, 0.0081]**<br>$p = 0.002$ |
| Native path only: temporal acquisition | **0.0091 [0.0085, 0.0098]**<br>$p = 0.016$ | **0.0110 [0.0068, 0.0153]**<br>$p = 0.002$ | **0.0223 [0.0214, 0.0233]**<br>$p = 0.016$ | **0.0298 [0.0161, 0.0436]**<br>$p = 0.002$ |
| Trilinear path only: spatial | **0.0023 [0.0012, 0.0034]**<br>$p = 0.016$ | **0.0023 [0.0013, 0.0033]**<br>$p = 0.002$ | **0.0041 [0.0020, 0.0063]**<br>$p = 0.016$ | **0.0033 [0.0016, 0.0050]**<br>$p = 0.002$ |
| Trilinear path only: temporal acquisition | **0.0091 [0.0085, 0.0098]**<br>$p = 0.016$ | **0.0110 [0.0068, 0.0153]**<br>$p = 0.002$ | **0.0223 [0.0214, 0.0233]**<br>$p = 0.016$ | **0.0298 [0.0161, 0.0436]**<br>$p = 0.002$ |
| Volume-scaled noise: spatial | 0.0002 [−0.0012, 0.0015]<br>$p = 0.781$ | 0.0003 [−0.0008, 0.0014]<br>$p = 0.512$ | 0.0024 [−0.0001, 0.0048]<br>$p = 0.047^{\ddagger}$ | **0.0018 [0.0001, 0.0036]**<br>$p = 0.033$ |

**Table S13.** Sensitivity to the temporal recovery method: point estimates, primary / broad cohort. No direct comparison between contrasts was planned for the alternative methods.

| Recovery method: contrast | nRMSE | PI | Trough (frames) | Peak (frames) |
|---|---|---|---|---|
| Periodic-linear (primary): spatial | 0.0032 / 0.0032 | 0.0054 / 0.0044 | 0.04 / 0.03 | 0.14 / 0.15 |
| Periodic-linear (primary): temporal acquisition | 0.0091 / 0.0110 | 0.0223 / 0.0298 | 0.63 / 0.76 | 0.72 / 0.72 |
| PCHIP: spatial | 0.0032 / 0.0032 | 0.0054 / 0.0044 | 0.04 / 0.03 | 0.14 / 0.15 |
| PCHIP: temporal acquisition | 0.0036 / 0.0056 | 0.0225 / 0.0300 | 0.63 / 0.76 | 0.72 / 0.72 |
| Sinc/Fourier: spatial | 0.0032 / 0.0032 | 0.0054 / 0.0044 | 0.04 / 0.03 | 0.14 / 0.15 |
| Sinc/Fourier: temporal acquisition | 0.0021 / 0.0043 | 0.0090 / 0.0165 | 0.36 / 0.52 | −0.25 / −0.18 |
| Nearest sample: spatial | 0.0032 / 0.0032 | 0.0054 / 0.0044 | 0.04 / 0.03 | 0.14 / 0.15 |
| Nearest sample: temporal acquisition | 0.0416 / 0.0427 | 0.0251 / 0.0325 | 1.13 / 1.21 | 0.97 / 0.90 |

**Table S14.** Other sensitivity analyses: inlet sections only, drift-corrected reference, the broad cohort without the case with an inlet-ramp cycle, and the corner-aligned operator.

| Sensitivity analysis | nRMSE: primary | nRMSE: broad | PI: primary | PI: broad |
|---|---|---|---|---|
| Inlet only: spatial | **0.0037 [0.0013, 0.0061]** $p = 0.016$ | **0.0038 [0.0017, 0.0059]** $p = 0.002$ | **0.0067 [0.0014, 0.0120]** $p = 0.016$ | **0.0054 [0.0016, 0.0092]** $p = 0.004$ |
| Inlet only: temporal acquisition | **0.0095 [0.0090, 0.0099]** $p = 0.016$ | **0.0112 [0.0071, 0.0154]** $p = 0.002$ | **0.0225 [0.0216, 0.0234]** $p = 0.016$ | **0.0295 [0.0162, 0.0427]** $p = 0.002$ |
| Linear drift correction: spatial | **0.0040 [0.0026, 0.0054]** $p = 0.016$ | **0.0038 [0.0025, 0.0051]** $p = 0.002$ | **0.0058 [0.0027, 0.0088]** $p = 0.016$ | **0.0045 [0.0019, 0.0071]** $p = 0.006$ |
| Linear drift correction: temporal acquisition | **0.0075 [0.0071, 0.0080]** $p = 0.016$ | **0.0076 [0.0072, 0.0080]** $p = 0.002$ | **0.0174 [0.0163, 0.0185]** $p = 0.016$ | **0.0187 [0.0168, 0.0206]** $p = 0.002$ |
| Without G3a: spatial | – | **0.0030 [0.0022, 0.0037]** $p = 0.002$ | – | **0.0045 [0.0024, 0.0065]** $p = 0.002$ |
| Without G3a: temporal acquisition | – | **0.0096 [0.0085, 0.0106]** $p = 0.002$ | – | **0.0248 [0.0212, 0.0285]** $p = 0.002$ |
| Corner-aligned operator: spatial | **0.0041 [0.0030, 0.0053]** $p = 0.016$ | **0.0039 [0.0028, 0.0050]** $p = 0.002$ | **0.0069 [0.0043, 0.0095]** $p = 0.016$ | **0.0054 [0.0029, 0.0078]** $p = 0.002$ |
| Corner-aligned − block-centred operator | **0.0010 [0.0002, 0.0017]** $p = 0.016$ | **0.0007 [0.0001, 0.0013]** $p = 0.018$ | **0.0016 [0.0003, 0.0028]** $p = 0.062^{\ddagger}$ | 0.0010 [−0.0001, 0.0020] $p = 0.074$ |

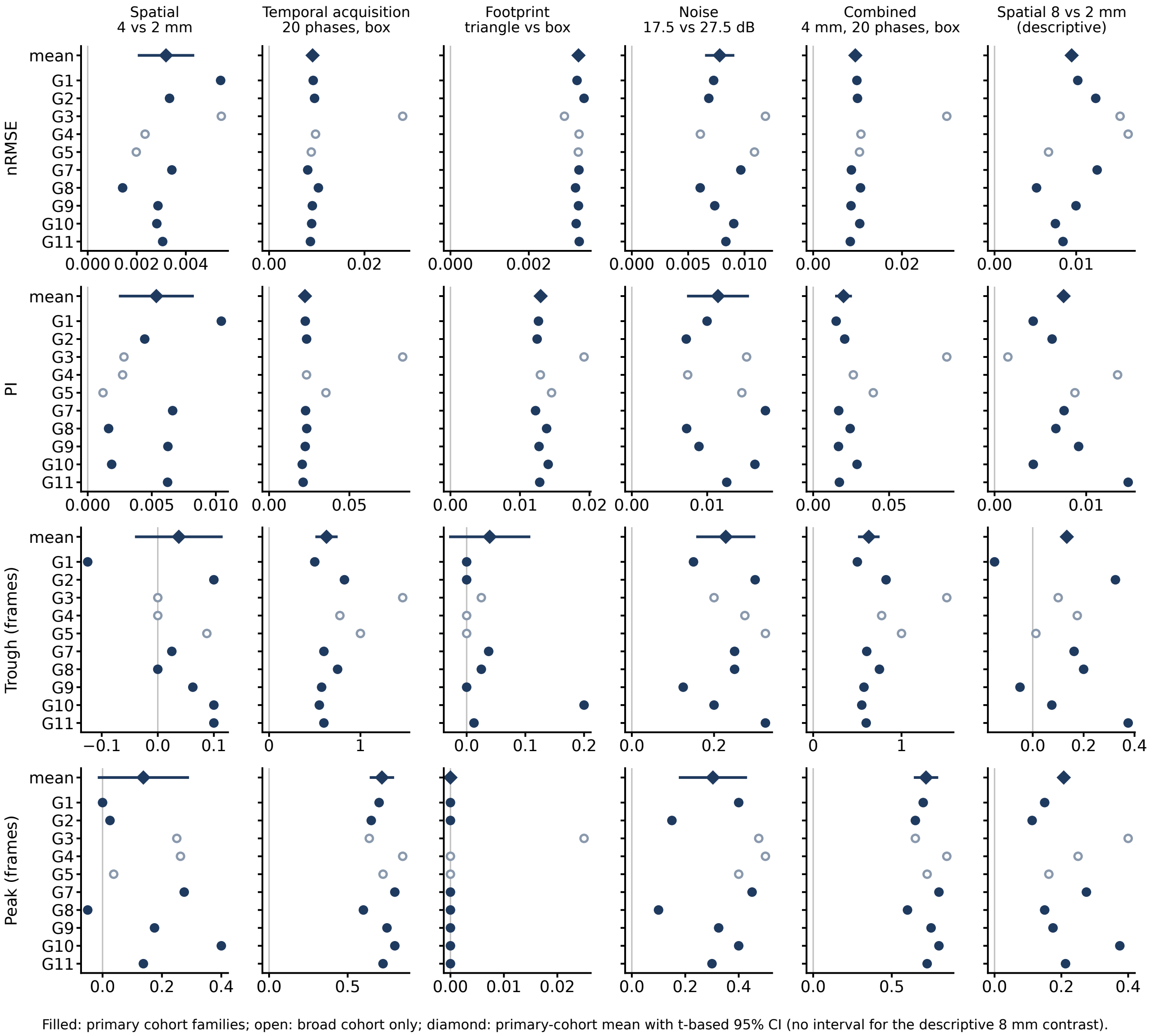


**Figure S1.** Family values of every contrast (columns) on the headline endpoints (rows). Filled circles: primary-cohort families; open circles: families in the broad cohort only; diamond: primary-cohort mean with t-based 95% CI (none for the descriptive 8 mm contrast).

**Table S15.** Headline contrasts and pre-specified direct comparisons, analysis v2.5.0 (periodic-linear recovery): t-based 95% CIs (primary inference) and family-bootstrap 95% CIs (secondary, for comparability with the earlier analysis). Primary: non-divergent cohort (7 families); broad: 10 families. Unadjusted for multiplicity. †The t-based interval includes 0 and the family-bootstrap interval excludes it. The t-based interval is the pre-specified inference rule (analysis plan, section 3); with few families the percentile bootstrap tends to give intervals that are too narrow, so these rows are not described as resolved.

| Contrast or comparison: endpoint | Primary: estimate | t 95% CI | bootstrap 95% CI | Broad: estimate | t 95% CI | bootstrap 95% CI |
|---|---|---|---|---|---|---|
| Spatial (4 vs 2 mm): nRMSE | 0.0032 | [0.0021, 0.0043] | [0.0024, 0.0041] | 0.0032 | [0.0023, 0.0042] | [0.0025, 0.0040] |
| Spatial (4 vs 2 mm): PI | 0.0054 | [0.0025, 0.0082] | [0.0034, 0.0075] | 0.0044 | [0.0023, 0.0065] | [0.0028, 0.0063] |
| Spatial (4 vs 2 mm): trough | 0.04 | [−0.04, 0.11] | [−0.02, 0.09] | 0.03 | [−0.02, 0.09] | [−0.01, 0.07] |
| Spatial (4 vs 2 mm): peak | 0.14 | [−0.01, 0.29] | [0.03, 0.25]† | 0.15 | [0.05, 0.26] | [0.07, 0.24] |
| Temporal acquisition (20 phases, box): nRMSE | 0.0091 | [0.0085, 0.0098] | [0.0087, 0.0096] | 0.0110 | [0.0068, 0.0153] | [0.0089, 0.0149] |
| Temporal acquisition (20 phases, box): PI | 0.0223 | [0.0214, 0.0233] | [0.0216, 0.0230] | 0.0298 | [0.0161, 0.0436] | [0.0222, 0.0422] |
| Temporal acquisition (20 phases, box): trough | 0.63 | [0.52, 0.74] | [0.56, 0.71] | 0.76 | [0.56, 0.97] | [0.62, 0.95] |
| Temporal acquisition (20 phases, box): peak | 0.72 | [0.65, 0.79] | [0.66, 0.77] | 0.72 | [0.67, 0.78] | [0.68, 0.77] |
| Footprint (triangle vs box): nRMSE | 0.0033 | [0.0032, 0.0033] | [0.0032, 0.0033] | 0.0032 | [0.0031, 0.0033] | [0.0031, 0.0033] |
| Footprint (triangle vs box): PI | 0.0130 | [0.0123, 0.0136] | [0.0125, 0.0135] | 0.0138 | [0.0123, 0.0152] | [0.0128, 0.0151] |
| Footprint (triangle vs box): trough | 0.04 | [−0.03, 0.11] | [0.01, 0.10]† | 0.03 | [−0.01, 0.07] | [0.00, 0.07]† |
| Footprint (triangle vs box): peak¶ | 0.000 | [0.000, 0.000] | [0.000, 0.000] | 0.002 | [−0.003, 0.008] | [0.000, 0.007] |
| Noise (17.5 vs 27.5 dB): nRMSE | 0.0078 | [0.0066, 0.0090] | [0.0070, 0.0087] | 0.0083 | [0.0069, 0.0098] | [0.0072, 0.0096] |
| Noise (17.5 vs 27.5 dB): PI | 0.0114 | [0.0075, 0.0154] | [0.0087, 0.0144] | 0.0117 | [0.0088, 0.0146] | [0.0094, 0.0142] |
| Noise (17.5 vs 27.5 dB): trough | 0.23 | [0.16, 0.30] | [0.18, 0.28] | 0.24 | [0.19, 0.29] | [0.20, 0.28] |
| Noise (17.5 vs 27.5 dB): peak | 0.30 | [0.18, 0.43] | [0.21, 0.39] | 0.35 | [0.25, 0.45] | [0.27, 0.42] |
| Combined (4 mm, 20 phases, box): nRMSE | 0.0095 | [0.0086, 0.0104] | [0.0088, 0.0102] | 0.0118 | [0.0071, 0.0164] | [0.0093, 0.0160] |
| Combined (4 mm, 20 phases, box): PI | 0.0200 | [0.0153, 0.0246] | [0.0169, 0.0237] | 0.0294 | [0.0137, 0.0451] | [0.0195, 0.0439] |
| Combined (4 mm, 20 phases, box): trough | 0.63 | [0.52, 0.74] | [0.56, 0.71] | 0.77 | [0.55, 0.99] | [0.62, 0.97] |
| Combined (4 mm, 20 phases, box): peak | 0.72 | [0.65, 0.79] | [0.66, 0.77] | 0.72 | [0.67, 0.78] | [0.68, 0.77] |
| nRMSE: spatial − temporal acquisition | −0.0059 | [−0.0074, −0.0045] | [−0.0071, −0.0049] | −0.0078 | [−0.0117, −0.0040] | [−0.0114, −0.0055] |
| PI: spatial − temporal acquisition | −0.0170 | [−0.0199, −0.0141] | [−0.0192, −0.0149] | −0.0254 | [−0.0399, −0.0110] | [−0.0388, −0.0167] |
| Trough timing: temporal acquisition − spatial | 0.59 | [0.48, 0.70] | [0.52, 0.67] | 0.73 | [0.52, 0.94] | [0.58, 0.92] |
| Peak timing: temporal acquisition − spatial | 0.58 | [0.49, 0.67] | [0.51, 0.64] | 0.57 | [0.50, 0.65] | [0.51, 0.63] |
| Mean-removed nRMSE: spatial − temporal acquisition | −0.0066 | [−0.0078, −0.0055] | [−0.0075, −0.0058] | −0.0087 | [−0.0129, −0.0045] | [−0.0126, −0.0063] |

¶Printed with 3 decimals (all other timing rows: 2) because the effect is below 0.005 frames in both cohorts.

**Table S16.** Family values of the spatial (Sp.) and temporal-acquisition (Temp.) contrasts on the headline endpoints, analysis v2.5.0 (primary cohort; values in parentheses and *: broad cohort only).

| Family | Sp. nRMSE | Sp. PI | Sp. trough | Sp. peak | Temp. nRMSE | Temp. PI | Temp. trough | Temp. peak |
|---|---|---|---|---|---|---|---|---|
| G1 | 0.0054 | 0.0104 | −0.12 | 0.00 | 0.0092 | 0.0225 | 0.50 | 0.70 |
| G2 | 0.0033 | 0.0044 | 0.10 | 0.02 | 0.0095 | 0.0234 | 0.82 | 0.65 |
| G3* | (0.0054) | (0.0028) | (0.00) | (0.25) | (0.0280) | (0.0832) | (1.46) | (0.64) |
| G4* | (0.0023) | (0.0027) | (0.00) | (0.26) | (0.0097) | (0.0233) | (0.78) | (0.85) |
| G5* | (0.0020) | (0.0012) | (0.09) | (0.04) | (0.0088) | (0.0354) | (1.00) | (0.72) |
| G7 | 0.0034 | 0.0066 | 0.02 | 0.28 | 0.0081 | 0.0227 | 0.60 | 0.80 |
| G8 | 0.0014 | 0.0016 | 0.00 | −0.05 | 0.0103 | 0.0234 | 0.75 | 0.60 |
| G9 | 0.0029 | 0.0063 | 0.06 | 0.18 | 0.0091 | 0.0225 | 0.57 | 0.75 |
| G10 | 0.0028 | 0.0019 | 0.10 | 0.40 | 0.0089 | 0.0206 | 0.55 | 0.80 |
| G11 | 0.0031 | 0.0062 | 0.10 | 0.14 | 0.0087 | 0.0212 | 0.60 | 0.72 |

**Table S17.** Source-data periodicity, analysis v2.5.0 (descriptive): cycle-closure (wrap) metric of the reference waveforms, mean over trunk sections, against the case values of the spatial and temporal-acquisition contrasts on nRMSE. *Broad cohort only.

| Case | Wrap metric (trunk mean) | Spatial nRMSE | Temporal nRMSE |
|---|---|---|---|
| G1b | 1.20 | 0.0054 | 0.0092 |
| G2a | 0.75 | 0.0032 | 0.0096 |
| G2b | 1.18 | 0.0034 | 0.0094 |
| G3a* | 5.10 | 0.0079 | 0.0426 |
| G3b* | 2.44 | 0.0030 | 0.0134 |
| G4a* | 1.07 | 0.0024 | 0.0097 |
| G4b* | 1.05 | 0.0023 | 0.0097 |
| G5a* | 1.26 | 0.0024 | 0.0097 |
| G5b* | 0.94 | 0.0015 | 0.0080 |
| G7a | 0.98 | 0.0031 | 0.0081 |
| G7b | 1.00 | 0.0037 | 0.0081 |
| G8a | 1.44 | 0.0014 | 0.0104 |
| G8b | 1.51 | 0.0014 | 0.0103 |
| G9a | 1.17 | 0.0030 | 0.0091 |
| G9b | 1.10 | 0.0028 | 0.0091 |
| G10a | 1.32 | 0.0028 | 0.0089 |
| G10b | 1.39 | 0.0028 | 0.0090 |
| G11a | 1.06 | 0.0031 | 0.0087 |
| G11b | 1.06 | 0.0030 | 0.0086 |
